# Within-host *Mycobacterium tuberculosis* evolution: a population genetics perspective

**DOI:** 10.1101/863894

**Authors:** Ana Y. Morales-Arce, Rebecca B. Harris, Anne C. Stone, Jeffrey D. Jensen

**Affiliations:** Center for Evolution and Medicine, Arizona State University, Tempe, Arizona, USA; School of Human Evolution and Social Change, Arizona State University, Tempe, Arizona, USA; School of Life Sciences, Arizona State University, Tempe, Arizona, USA

## Abstract

The within-host evolutionary dynamics of TB remain unclear, and underlying biological characteristics render standard population genetic approaches based upon the Wright-Fisher model largely inappropriate. In addition, the compact genome combined with an absence of recombination is expected to result in strong purifying selection effects. Thus, it is imperative to establish a biologically-relevant evolutionary framework incorporating these factors in order to enable an accurate study of this important human pathogen. Further, such a model is critical for inferring fundamental evolutionary parameters related to patient treatment, including mutation rates and the severity of infection bottlenecks. We here implement such a model and infer the underlying evolutionary parameters governing within-patient evolutionary dynamics. Results demonstrate that the progeny skew associated with the clonal nature of TB severely reduces genetic diversity and that the neglect of this parameter in previous studies has led to significant mis-inference of mutation rates. As such, our results suggest an underlying *de novo* mutation rate that is considerably faster than previously inferred, and a progeny distribution differing significantly from Wright-Fisher assumptions. This inference largely reconciles the seemingly contradictory observations of both rapid drug-resistance evolution but extremely low levels of genetic variation in both resistant and non-resistant populations.

## INTRODUCTION

Tuberculosis (TB) is a public health threat worldwide (WHO, 2018). Despite clear motivation for study, the observed within- and between-host evolutionary dynamics of *Mycobacterium tuberculosis* (*M.TB*) are not well understood, and results to date represent something of a paradox. On the one hand, drug resistance evolves rapidly (Fonseca *et al*. 2015; Eldholm *et al*. 2015); on the other, the genomic characteristics of *M.TB* do not appear conducive for such rapid adaptation, with inferred mutation rates being amongst the slowest of any human pathogen (Rocha *et al*. 2006; Ford *et al*. 2011; Ford *et al*. 2013; Colangeli *et al*. 2014; Payne *et al*. 2019; Menardo *et al*. 2019) and remarkably little genetic variation observed within or between hosts. Furthermore, purifying selection has been argued to play both a dominant as well as a weak role in shaping patterns of variation (Hershberg *et al*. 2008; Pepperell *et al*. 2013), and demographic estimates suggest a population history of TB that either matches or is uncorrelated with that of its human host (Comas *et al*. 2013; Bos *et al*. 2014; Brites *et al*. 2015; Eldholm *et al*. 2016).

To obtain a more robust understanding of TB evolutionary dynamics, it is essential to first appreciate that between-population observations are simply an aggregation of within-population processes. As such, studying the population genetics of within-patient data is critical to understanding the genetic differences observed between patients as well as their treatment outcomes. Fortunately, recent advances in sequencing technologies have allowed for more abundant and higher quality within-patient data. These published datasets have revealed a few common features of *M.TB*, including low-levels of genome-wide variation. For instance, Trauner *et al*. (2017) deep-sequenced twelve patients across four-time points and observed fewer than 50 polymorphic sites per patient genome-wide. In addition, the observed site frequency spectrum (SFS) is generally characterized by an abundance of rare variants (*i.e*., it is strongly left-skewed). These patterns have partly led to the suggestion that purifying selection effects may be wide-spread in the *M.TB* genome (Brown *et al*. 2016; Phelan *et al*. 2016; Mortimer *et al*. 2018).

Additional evolutionary factors likely contribute to these genomic patterns as well. For example, population bottlenecks may reduce genetic variation and alter the shape of the SFS (see review Thornton *et al*. 2007). Previous *M.TB* studies have investigated these effects separately in both the deep-time view of the population bottleneck and subsequent growth experienced by the host human population (Hershberg *et al*. 2008, Liu *et al*. 2018), as well as the shallow-time view of the population bottleneck and subsequent growth characterizing each novel transmission event and treatment (*e.g.,* Trauner *et al*. 2017). Additionally, in fitting the left-skewed SFS, Pepperell *et al*. (2013) found that such a demographic history combined with a mix of both deleterious and neutrally-evolving sites produced the nearest fit to the observed SFS. Finally, given the lack of recombination in *M.TB*, related linkage effects (*i.e*., background selection (Charlesworth *et al*. 1993)) have similarly been discussed within these contexts (Pepperell *et al*. 2010; Copin *et al*. 2016).

While these studies have provided many important insights, there remains a relatively unexplored, though potentially highly significant, effect: clonality. Indeed, clonality and the related progeny distribution represents an important violation of commonly used evolutionary inference approaches based upon the Wright-Fisher (WF) model and the related Kingman coalescent (Eldon *et al*. 2006; Dos Vultos *et al*. 2008; Huillet *et al*. 2011; Lapierre *et al*. 2016). Specifically, progeny distributions under the WF model are Poisson distributed with a mean and variance of 1. Therefore, when an individual produces many offspring, far in excess of simple replacement in the next generation, the assumption that only two lineages coalesce at a time is violated, resulting in multiple-merger coalescent (MMC) events (see reviews of Tellier & Lemaire 2014; Irwin *et al*. 2016).

While perhaps abstract at first blush, this violation has very important implications for the study of sequence variation and diversity. Namely, as *M.TB* has been found to exhibit strong progeny skew owing to obligate clonal reproduction (Baker *et al*. 2004; Dos Vultos *et al*. 2008), the null model against which the above studies are comparing becomes incorrect. For example, under a multiple-merger model, the effective population size (*N_e_*) no longer scales linearly with census size (*N*) as it does under the Kingman coalescent (Huillet *et al*. 2011). As a result, genetic diversity is a nonlinear function of the underlying population size - a result of interest given the strongly constrained and similar levels of variation observed across TB patients, regardless of infection time or resistance status. Similarly, under these progeny-skew models, the SFS is skewed towards an excess of low-frequency variants, generating a negative Tajima’s *D* even under equilibrium neutrality (Eldon *et al*. 2006; Birkner *et al*. 2013; Blath *et al*. 2016) – which appears of relevance to *M.TB* populations given the pervasively left-skewed SFS observed both within and between TB patients. Finally, the fixation probability of beneficial mutations under progeny skew may become much larger than under the WF model, owing to the increased probability of rapidly escaping stochastic loss (Der *et al*. 2011). This is fundamental to understanding the rapidly and independently evolving drug-resistance mutations in global TB populations - a result seemingly at odds with the previously inferred mutation rates (Sherman *et al*. 2011; Colangeli *et al*. 2014; Duchêne *et al*. 2016). In sum, the general theoretical expectations owing to progeny skew alone appear to qualitatively match empirical observations from *M.TB*; observations which, to date, have been attributed to alternate processes.

Recent progress has been made in utilizing these models to disentangle and even co-estimate patterns of demography, progeny skew, and selection. While there exist a variety of potential MMC models (see review of Tellier & Lemaire 2014), the so-called *Ψ*-coalescent has been a major focus of this literature given the straight-forward biological interpretation. Namely, the parameter *Ψ* represents the proportion of the next generation arising from a single parent (*e.g., Ψ* = 0.05 implies that one individual contributes offspring that comprise 5% of the next generation). In addition to which, recent experimental measures from viral populations are offering real-time insights in to such progeny distributions (Vahey and Fletcher 2019). Three results are of particular importance here. First, Eldon *et al*. (2015) demonstrated that population growth may be distinguished from multiple-merger coalescent events owing to progeny-skew, given differing expectations in the SFS. Second, Matuszewski *et al*. (2018) derived analytical expectations for the SFS under a multiple merger coalescent model with changing population size and further demonstrated that these parameters can indeed be accurately inferred jointly within a likelihood framework. Finally, building upon the two above results as well as the approximate Bayesian statistical framework developed by Foll *et al*. (2014, 2015), Sackman *et al*. (2019) recently extended these results and demonstrated an ability to co-estimate progeny skew, effective population size, as well as per-site selection coefficients from time-sampled polymorphism data.

Thus, a tremendous opportunity now exists to understand better the impact of mutation, genetic drift, and selection in dictating patterns of *M.TB* evolution and genomic variation under this more realistic coalescent model accounting for the underlying progeny distributions inherent to clonal reproduction. While the approximate Bayesian approach of Sackman *et al*. (2019) would appear ideal for this purpose, it is a time-sampled estimator reliant upon considerable levels of segregating variation in order to track changing allele frequencies (as is commonly observed in viral populations, for example). Thus, this approach is under-powered given the minimal levels of variation observed in *M.TB*. Further, as the underlying mutation rate itself is a question of great interest and importance in *M.TB* (Payne *et al*. 2019), it is desirable to additionally co-estimate this parameter rather than assume it to be known.

With this motivation, we developed a novel statistical approach utilizing the insights described above pertaining to infection dynamics and widespread purifying selection, while overlaying inference of underlying mutation rates, progeny distributions, and the severities of infection-related population bottlenecks. By re-analyzing published data within this framework, our results demonstrate the great importance of the multiple merger consideration and resolve many of the evolutionary paradoxes surrounding *M.TB*.

## MATERIALS AND METHODS

### Simulations

We conducted forward-in-time simulations using the SLIM version 3 software package (Haller & Messer 2019). *M.TB* populations were modeled using a genome size of 441,153 kb, equivalent to a tenth portion of the true genome size, for computational efficiency. As *M.TB* is a compact, highly-coding genome (Cole *et al*. 1998; Fleischmann *et al*. 2002), we assumed a distribution of fitness effects (DFE) characterized equally by deleterious (*s* = −0.01) and nearly neutral (*s*=-0.001) mutations. Mutation rate (*μ*) measured *in vitro* has been reported to be as slow as 2e-10 (Ford *et al*. 2011). In contrast, higher estimates ranging from 1e-9 to 9e-6 (Ford *et al*. 2013) have been proposed; therefore, our study considered the full extent of this range. Furthermore, it is important to note that previous experiments have measured only the neutral mutation rate, not the total mutation rate. In other words, the large input of strongly deleterious mutations - comprising a substantial component of the total mutation rate - has not been included in earlier estimates as these mutations are unlikely to be sampled as segregating variation or as fixed differences. However, as these mutations are important for shaping diversity via both purifying selection and background selection effects, and as our interest is in understanding the total rate at which all *de novo* mutations are input into the population, we considered the total rather than the neutral mutation rate. In order to infer this parameter within the context of an appropriate progeny-skew model, *μ* was drawn from a prior uniform distribution between 1e-9 and 9e-6 per site per generation.

Following an initial burn-in period of 10*N* generations, we considered a three-stage demographic model characterizing a single patient infection: moving forward in time, we describe 1) a neutral equilibrium population of size *N*, 2) an initial infection bottleneck leading to an instantaneous population reduction to size *N2*, and 3) a subsequent population size recovery to size *N*. In stage 1, we modeled a population of size *N* = 1,000. In order to quantify the effects of underlying assumptions pertaining to population size, additional simulations and inference were performed at *N* = 25,000. During stage 2, the severity of the population bottleneck (*β*) was sampled from ∼U[0.001, 0.1], where *N2* = *N*β* - as the distribution of infection size in humans is unknown. However, it has been reported that in cattle TB (*M. bovis*) infection can be established by a single cell forming unit (Dean *et al*. 2005). During stage 2, the degree of progeny skew (or *Ψ*) was sampled from a prior distribution of ∼U[0, 0.2]. A value of 0 corresponds to the standard WF model. Progeny skew was simulated following the procedure of Sackman *et al*. 2019. In brief, one individual is chosen from the primary population A and founds a separate subpopulation B, the single generation unidirectional migration rate from B to A is set to *Ψ*, and the chosen individual thus contributes *NΨ* offspring to the following generation of A. A series of mate choice callbacks in SLiM force the migration rate to be exact rather than stochastic (see Supplementary Materials of Sackman *et al*. 2019). Subpopulation B is removed, the next generation begins, and a new individual is randomly sampled for the following generation. As such, each generation is a combination of *N*(1-*Ψ*) replacement events and a single sweepstakes event of magnitude *NΨ.* To emulate patient sampling at the onset of symptoms (approximately three months minimum (Behr *et al*. 2018)), we allowed stage 2 to run for 90 generations, assuming a generation time of 24 hours (Cole *et al*. 1998), and stage 3 to run for 910 generations before outputting genome alignments in ms format. Thus, the total generation time of our model was 11*N*.

Drawing from these prior distributions, 10,000 points (*i.e.,* parameter combinations) were sampled. For each parameter combination, we conducted 1,000 replicates in order to characterize both the mean and variance. Summary statistics were calculated in the R package PopGenome version 2.6.1 (Pfeifer *et al*. 2014).

### Data analysis and joint posterior estimates

For comparison to patient data, we examined the distribution and mean of segregating sites in samples published by Trauner *et al*. (2017). (https://zenodo.org/record/322377#.XO2CAy2ZNBw). A subset of 1,000 populations was simulated under the conditions described above. Given that patient data are subject to stringent filtering criteria, removing SNPs under a particular frequency cut-off, it was necessary to filter the simulated data to allow fair comparison. Thus, 100 genomes were sampled per each simulated population, and variants < 2% frequency were filtered out before the calculation of summary statistics (Table S1). Afterward, only genomes with ≥ 2 segregating sites were considered for further analyses, and the fraction of “invariable genomes” was noted for comparison to patient samples (Figure 2 and Table S1).

## RESULTS & DISCUSSION

### Considering levels of variation

We here report the first joint consideration of mutation rate, purifying selection, infection history, and progeny-skew in *M.TB* populations. A correlation of summary statistics (Figure 1) demonstrates that while mutation rate (*μ*) increases the number of segregating sites as expected, progeny skew (*Ψ*) acts to reduce variation. Furthermore, as the *Ψ*-parameter is of relevance every generation (*i.e*., every reproduction event), the impact of this previously unconsidered progeny skew on levels of variation is in fact much stronger than the single bottleneck event associated with infection. Considering the full distribution of sampled *μ* values (Figure S1), it is apparent that *Ψ* drastically reduces the average proportion of segregating sites genome-wide even for fast mutation rates, and that the probability of producing genomes devoid of any variation will naturally increase as *μ* decreases.

**Figure 1.**
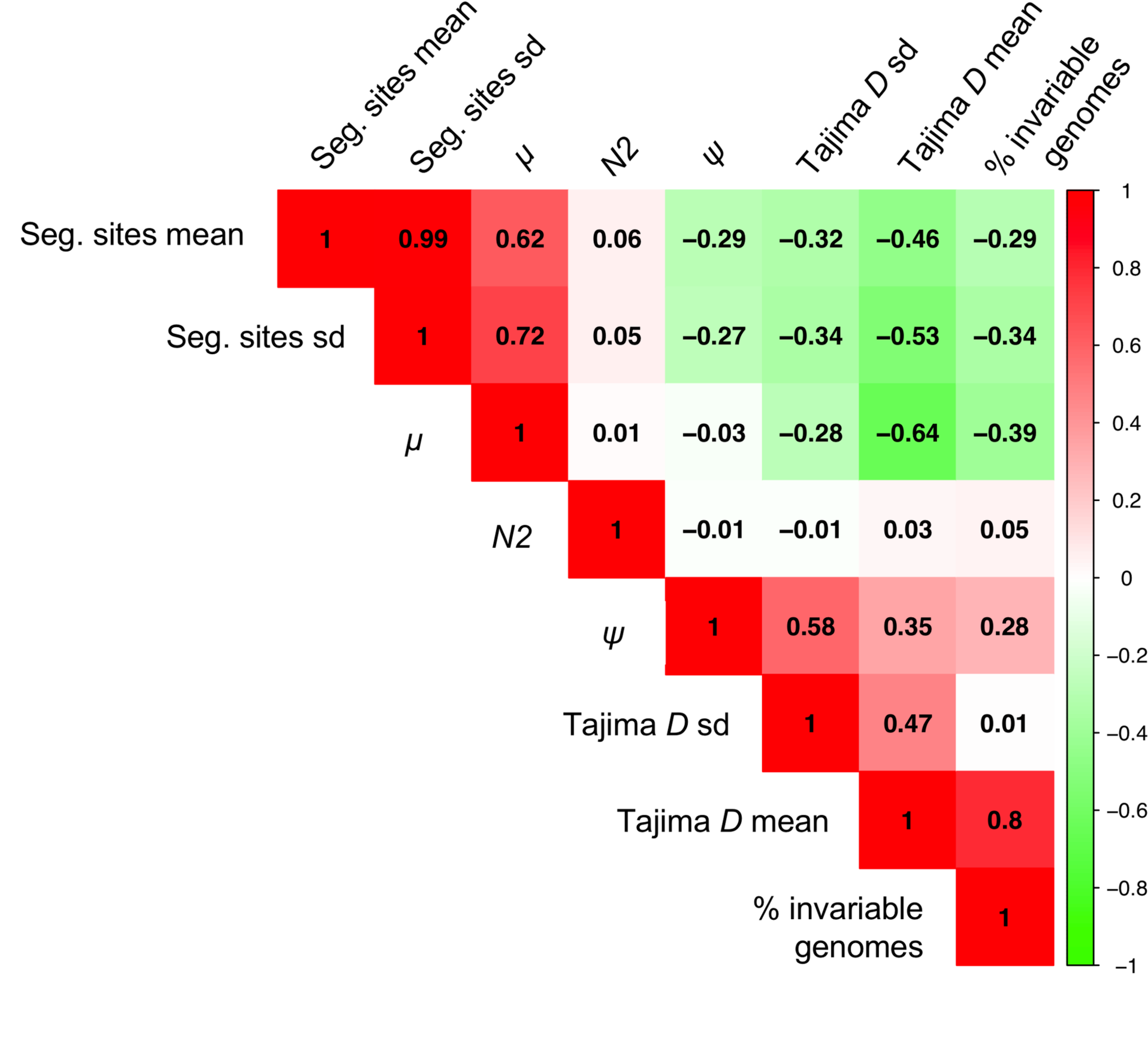
Correlation heatmap of parameters and summary statistics. Correlations are given between the parameters of interest (mutation rate (*μ*), progeny skew (*Ψ*), and bottleneck severity (*N2*)), and summary statistics (the mean and variance of the level of variation as measured by the number of segregating sites, and the mean and variance of the distribution of variation as measured by Tajima’s *D*). As shown, *N2* values do not correlate with any of the summary statistics, as the effect of the single generation bottleneck is swamped by the per-generation reproductive skew. Further, as expected, values of *μ* positively correlate with the number of segregating sites, while *Ψ* acts to reduce variation and is thus negatively correlated. Finally, while *μ* would not be expected to strongly correlate with the shape of the SFS (here summarized by Tajima’s *D*) for neutral mutations, it does so here given that we explicitly account for the input of deleterious mutations (see Methods).

In order to consider a range of *μ* consistent with observed data - once pervasive purifying selection and progeny skew have been taken in to account - two examples of patient data collected by Trauner *et al*. (2017) representing low (20 segregating sites genome-wide) and high (50 segregating sites genome-wide) variation samples were plotted and compared to the simulated data. In order to compare simulated data with patient data, the same filtering steps must be applied. In this case, SNPs under 2% frequency were filtered in the empirical data, and thus, the simulated data were similarly filtered in order to be comparable (Figure 2).

**Figure 2.**
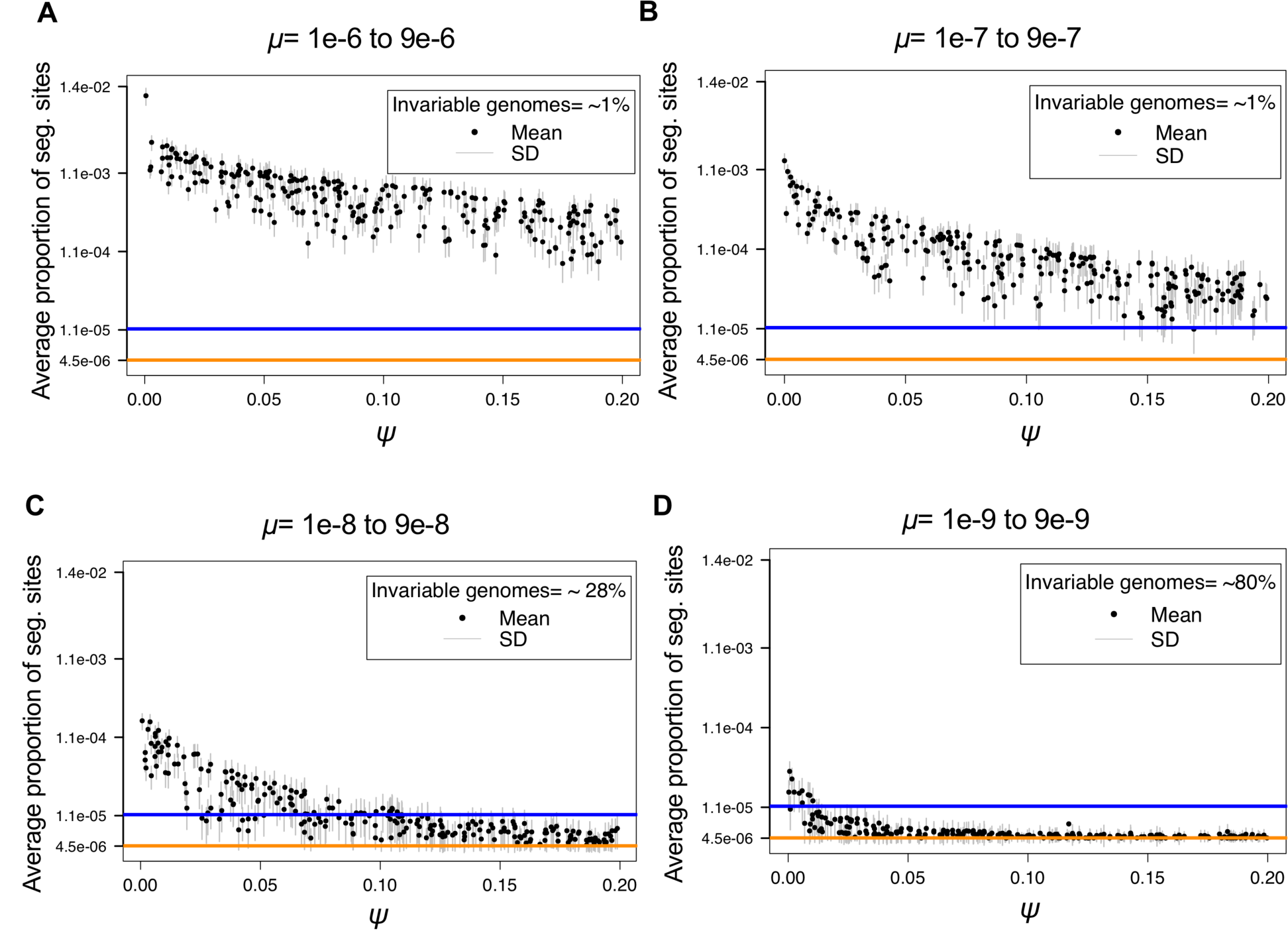
Log scale distribution of segregating sites above 2% frequency, as a function of mutation rate (*μ*) and progeny skew (*Ψ*). For each parameter combination (1,000 in total) of *μ* and *Ψ* drawn from the prior distributions, 1,000 replicates were simulated, with the mean given by the black dot and the standard deviation given by the gray bars. Each panel corresponds to a different order of magnitude of mutation rate range: (Α) 1e-6 to 9e-6, (B) 1e-7 to 9e-7, (C) 1e-8 to 9e-8, and (D) 1e-9 to 9e-9. The colored lines correspond to two examples of the proportion of segregating sites observed genome-wide in empirical patient data: 20 segregating sites as a mean (orange), and 50 segregating sites from patient_10 (blue) (Trauner *et al*. 2017). As shown, the range of segregating sites for the fast mutation rates (panels A and B), result in expectations much larger than that observed in patient data, regardless of *Ψ*. Conversely, the slowest mutation rate (panel D), result in too little variation, except under WF conditions (*i.e*., *Ψ* near 0) which are known to be violated given clonality. Thus, rates on the order of 1e-8 to 9e-8 (panel C) appear to well explain the range of variation observed in patient data, and further imply values of *Ψ* ranging roughly from 0.05 to 0.1, consistent with values previously estimated for within-host virus data (Sackman *et al*. 2019).

As shown, fast mutation rates (*μ* on the order of 1e-6 and 1e-7) routinely produce expected numbers of segregating sites far above that observed in patients, regardless of other parameter values, while slow mutation rates (*μ* on the order of 1e-9) generally result in too little variation to match observation. This result is of interest as *M.TB* mutation rates are generally believed to be exceedingly slow (Sherman *et al*. 2011) - though importantly, this inference has largely neglected the important contribution of these additional evolutionary processes. Thus, once accounting for the diversity-reducing effects inherent to clonality, as well as the extent of purifying selection effects inherent to a compact, non-recombining genome, it is evident that the *de novo* mutation rate is likely faster than previously believed, with mutation rates on the order 1e-8 well matching the range of observed data (Figure 2). In addition, in order to consider the impact of underlying assumptions pertaining to population size, simulations were re-performed with a 25x larger population size. As shown in Figure S2, owing to these diversity reducing effects, mutations rates on the order of 1e-8 remain the best fit to observed levels of variation, with slower mutation rates still producing too little variation and too many invariable genomes to be consistent with patient data.

### Considering distributions of variation

For the general range of *μ* identified above, the number of genome-wide segregating sites in simulated population data ranged from a minimum average of 2.1 to a maximum average of 78.7 SNPs (Table S1), depending on the combination of *μ* and *Ψ* drawn from the priors. Specifically, higher values of *μ* may be off-set by higher values of *Ψ*, resulting in a similar number of segregating sites for multiple parameter combinations. For example, for the patient sample containing 10 SNPs, simulation results demonstrate that *μ* = 8.13e-08 and a *Ψ* =0.06 would produce an average of 10.15 +/-4.64 SNPs, potentially suggesting a good fit to the data. However, *μ* = 3.1e-08 and a *Ψ* =0.02 can also generate similar results, yielding 10.40 +/-4.45 SNPs. More generally, this ridge in the posterior distributions (Figure 3A) between these two parameters suggests that they will be difficult to estimate independently if only levels of variation are used.

**Figure 3.**
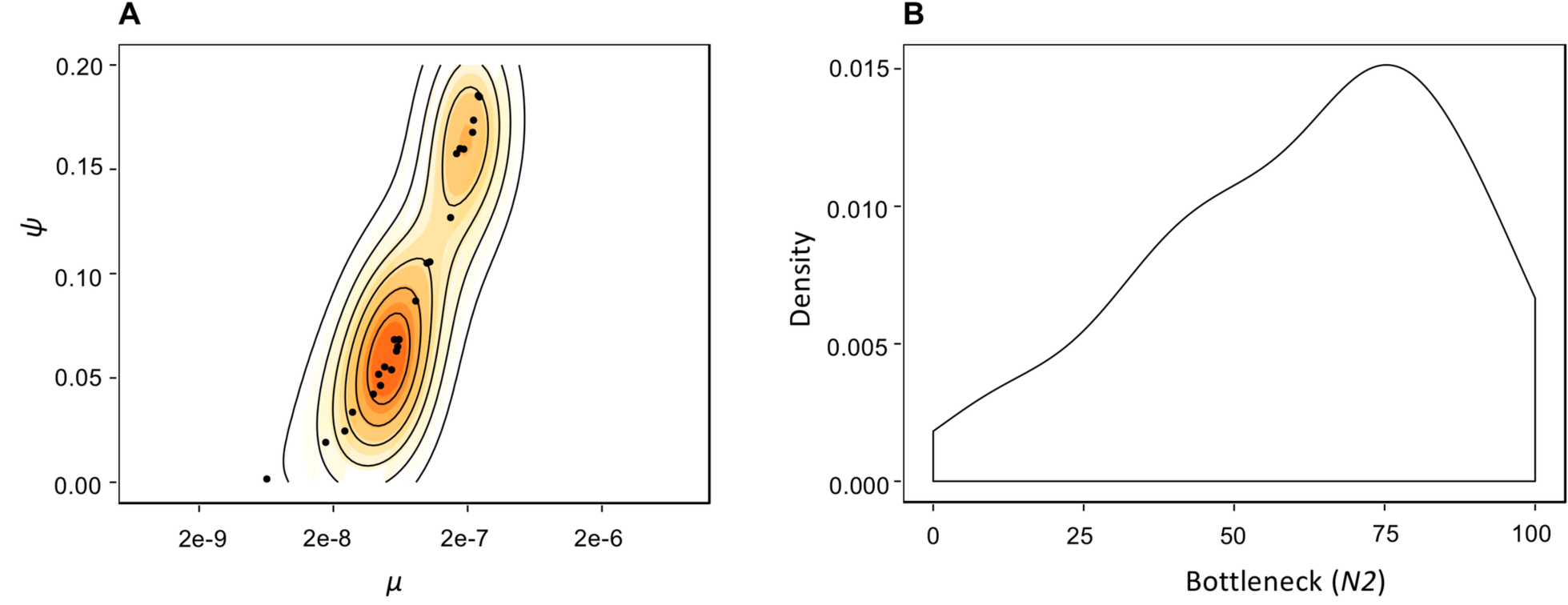
Posterior parameter estimates pertaining to patient data. (A) Joint posterior distribution for the parameters *μ* and *Ψ*. Solid contour lines specify the highest posterior density intervals. Owing to the diversity-increasing effect of *μ* and diversity-decreasing effect of *Ψ*, there exists a ridge in the joint posterior. Regardless, owing to differing expectations in the SFS along this ridge, inference suggests *μ* values within the 1e-8 range in combination with *Ψ* < 0.10. (Β) Posterior density for the severity of the infection bottleneck (*N2*). The X-axis gives the number of genomes at the time of infection reduced from 1,000. While the posterior distribution is non-uniform, the observation that all tested values remain consistent with patient data strongly suggests that there is not sufficient information in the data to estimate this third parameter (*i.e.,* size of the bottleneck, in addition to *μ* and *Ψ*).

Thus, while comparisons with general levels of variation are useful for identifying a range as in the above section, more information is needed to parse values further. Importantly, previous theoretical results (Eldon & Wakeley 2016; Matuszewski *et al*. 2018) have well described the effect of *Ψ* on the observed distribution of genetic variation (*i.e*., SFS). To utilize this information, a general summary of the SFS, Tajima’s *D* (1989), was calculated on the filtered simulated data. As shown (Figure S3), the shape of the SFS, and thus the value of the *D-* statistic, is related to the value of *Ψ*. As the degree of progeny skew initially increases, *D* becomes increasingly negative, as previously described. However, as progeny distributions become highly skewed, levels of variation are sharply reduced, resulting in an apparent increase in *D* values (Figure S3). Increasing *D* values could also be a result of the underlying filtering criteria, as after filtering simulations to match real data with segregating sites > 2%, *D* values increased proportionally (Figure S4). However, Tajima’s *D* is consistently negative in all mutation ranges, even after filtering.

### Parameter inference from patient samples

Thus, we next considered these results in light of published patient data. Recent publications have suggested that NGS technologies could facilitate personalized treatment in TB patients, allowing for improved outcomes (Copin *et al*. 2016; Cancino-Muñoz *et al*. 2019). To utilize such data, however, it is vital to understand the evolutionary dynamics shaping within-host *M.TB* diversity. As an illustrative example, we have re-examined the number of segregating sites in patient samples and estimated a mean ∼10 segregating sites per sample (Trauner *et al*. 2017) (Figure S5). Further, using the results and expectations obtained above regarding the level and distribution of variation, the patient data appear best fit by simulated populations with *μ* values ranging from 7.3e-9 to 3.8e-7, with the strongest posterior density at *μ ∼* 6e-8 and *Ψ* ∼0.06 (Figure 3A). Notably, faster *μ* values could produce similar results, but only if *Ψ* proportions are in excess of 0.15 (Figure 3A).

In addition, owing to the strong per-generation reductions associated with progeny skew, there remains no signal in the data to estimate the severity of the infection bottleneck accurately (Figure 3B). Specifically, while there is an increase in density towards stronger bottleneck values, the posterior distribution is not notably distinct from the prior distribution ∼U[0.001, 0.1] (Figure 3B). Apart from the population size reduction associated with infection, these results have important implications for the ability to detect other parameters of clinical interest - namely, the presence of selective sweeps associated with beneficial mutations (*e.g.,* potentially owing to drug-resistance). First, while there is strong statistical power to infer both *μ* and *Ψ* from patient data, there is little power to detect isolated events in the past (Figure 3B). This result, though unexpected under standard WF assumptions, is intuitive given the non-WF progeny distributions related to clonality. Namely, the diversity reduction associated with a single bottleneck event multiple generations in the past is not discernible from the per-generation diversity reduction related to clonal reproduction. Further, given that a selective sweep is, in fact, a type of population bottleneck (Barton 1998), this result also demonstrates that detecting selective sweeps associated with resistance mutations in this non-recombining organism, based on levels and patterns of genomic variation, will be exceedingly difficult. However, this observation well reconciles the fact that levels of variation do not appear significantly different between resistant and non-resistant *M.TB* patient populations (Trauner *et al*. 2017) - that is, these additional evolutionary processes are shaping variation so strongly, that the presence or absence of a resistance-associated selective sweep does not result in strongly differentiable expectations.

### Implications for characterizing the history of TB in humans

A topic of wide-spread interest in the literature pertains to the history of TB in the human host. This inference has primarily been made within a phylogenetic context, relying on the construction of a single consensus sequence per patient. While such a comparison of consensus sequences can be highly mis-leading when making evolutionary inference (see Renzette *et al*. 2017), these age-estimates also inherently rely on an accurate knowledge of mutation rates in order to invoke the ‘clock-like’ accumulation of neutral mutations as a proxy for time (Menardo *et al*. 2019). As our results demonstrate that previous mutation rate estimates have likely been downwardly-biased, it is of interest to consider what these revised mutation rates would imply for this evolutionary history. However, there are at least three difficulties in directly comparing population-level estimates with previous consensus-based phylogenetic inference. Firstly, estimates are generally given per year, whereas the preferred evolutionary rate is per generation (as given here). There is support for one generation per day as a conversion (Cole *et al*. 1998), though further study is necessary to quantify the correct scaling factor. Secondly, when invoking a divergence-based clock, previous studies are measuring the neutral mutation rate, given that the rate of mutation is equivalent to the rate of divergence for neutral mutations only (Kimura 1968). However, we are here interested in the total mutation rate (that is, the rate at which neutral and non-neutral mutations arise per generation); therefore, our rate must be parsed into neutral and non-neutral components to enable appropriate comparison. Similar to the first point, additional research is necessary in order to better quantify the distribution of fitness effects in *M.TB,* as understanding the fraction of total mutations represented by neutral mutations is necessary for the conversion. Finally, we here consider the alleles segregating within a population for inference (*i.e.,* within a patient), whereas previous studies often call a consensus sequence per patient (*i.e.,* per population) and make inferences based on a collection of such consensus sequences. Such a summary of population-level variation into a single sequence is difficult to interpret, though what is evident is that a great majority of rare alleles will be neglected, and thus only a small subset of total variation (*i.e.,* common alleles) will be considered (Renzette *et al*. 2017). As such, we propose that future evolutionary inference pertaining to TB would benefit tremendously from a full consideration of within-patient diversity, as we demonstrate here. In sum, any direct comparison with consensus-based phylogenetic age estimates would be overly speculative at this juncture.

## CONCLUSIONS

TB patient infection dynamics have remained enigmatic. We here argue that much of the difficulty in interpreting patterns of variation and evolution has owed to an inappropriate underlying null model, relying on classical expectations developed for organisms with very different underlying biological properties. Fortunately, recent theoretical extensions in non-WF and alternative coalescent models, more appropriate for clonally reproducing organisms, have created an opportunity to revisit existing *M.TB* patient data. By accounting for the pervasive purifying selection effects associated with this non-recombining, highly-coding genome, as well as the skewed progeny distributions inherent to clonal reproduction, we have provided improved insights into the evolutionary dynamics shaping within-host variation. Further, through a consideration of these diversity-reducing effects, results suggest an underlying *de novo* mutation rate that is considerably faster than previously inferred. This largely reconciles the seemingly contradictory observations of both rapid resistance evolution, but extremely low levels of population variation. Namely, the population mutation rate may indeed be sufficiently fast to provide a steady input of beneficial mutations, explaining the rapid resistance evolution clinically observed. However, recurrent purifying selection and progeny skew act together to rapidly eliminate segregating variation from the population, reconciling the minimal levels of variation observed as well as the general homogeneity in levels and distributions of variation in both resistant and non-resistant patient samples alike. Furthermore, the role of these per-generation evolutionary forces in shaping patterns of variation is sufficiently strong that periodic events, including the infection-associated bottleneck and selective sweeps centered on drug-resistance mutations, will be challenging to detect and quantify on top of these more common processes.

## Supporting Information

**Figure S1.**
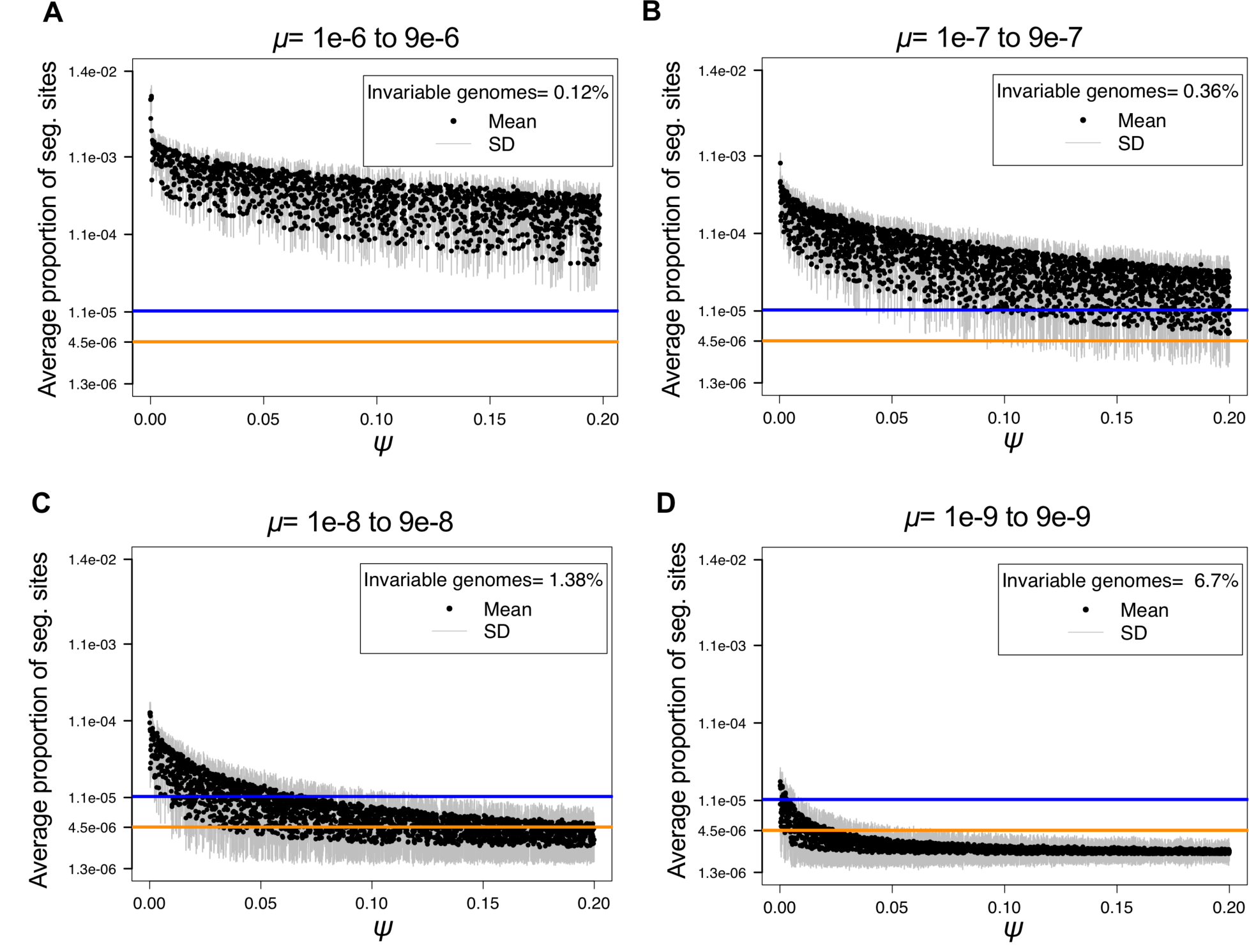
Log scale distribution of segregating sites in unfiltered simulated populations as a function of mutation rate (*μ*) and progeny skew (*Ψ*). For each parameter combination of *μ* and *Ψ* drawn from the prior (10,000 in total), 1,000 replicates were simulated, with the mean given by the black dot and the standard deviation given by the gray bars. *Ψ* proportions are given on the X-axis, while the proportion of segregating sites observed in the genome are given on the Y-axis. Each panel corresponds to a different order of magnitude of mutation rate: (Α) 1e-6 to 9e-6, (B) 1e-7 to 9e-7, (C) 1e-8 to 9e-8, and (D) 1e-9 to 9e-9. The colored lines correspond to two examples of the proportion of segregating sites observed genome-wide in empirical patient data: 20 segregating sites as a mean (orange) and 50 segregating sites as the maximum observed from patient_10 (blue) (Trauner *et al*. 2017).

**Figure S2.**
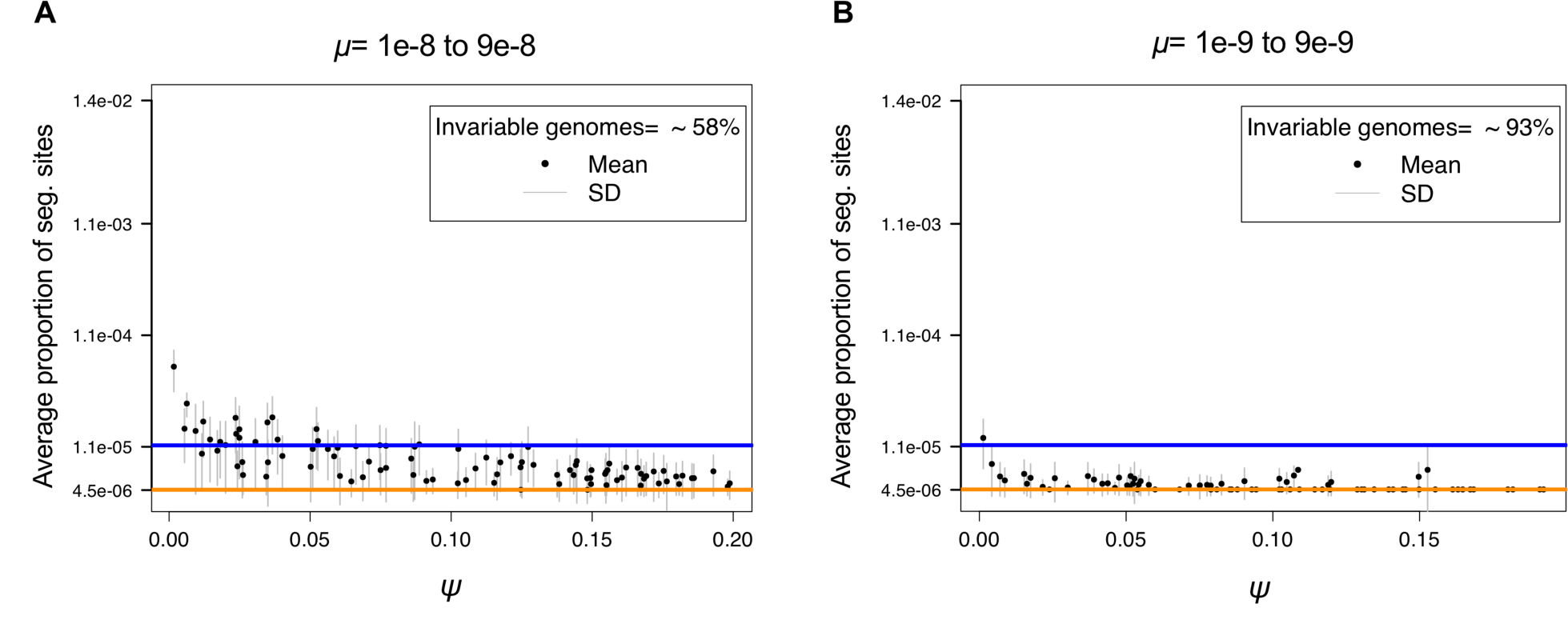
Log scale distribution of segregating sites in filtered simulated populations of *N*= 25,000 as a function of mutation rate (*μ*) and progeny skew (*Ψ*). For each parameter combination of *μ* and *Ψ* drawn from the prior (100 in total), 100 replicates were simulated, with the mean given by the black dot and the standard deviation given by the gray bars. *Ψ* proportions are given on the X-axis, while the proportion of segregating sites observed in the genome are given on the Y-axis. Each panel corresponds to a different order of magnitude of mutation rate: (Α) 1e-8 to 9e-8, and (B) 1e-9 to 9e-9. The colored lines correspond to two examples of the proportion of segregating sites observed genome-wide in empirical patient data: 20 segregating sites (orange) and 50 segregating sites (blue) (Trauner *et al*. 2017). As shown, inference remains consistent with the results of **Figure 2**, even under this 25x larger population size.

**Figure S3.**
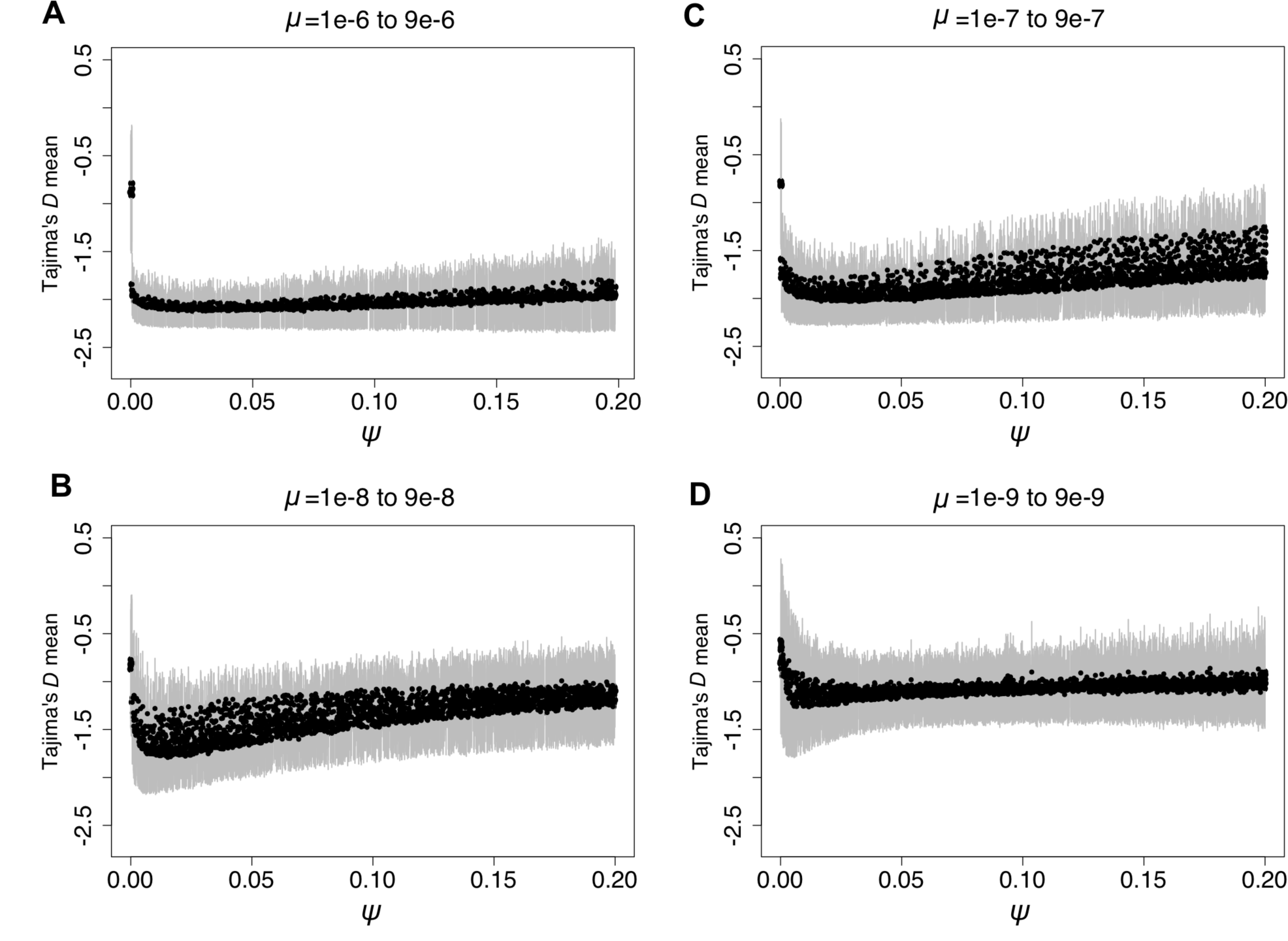
The shape of the site frequency spectrum in unfiltered simulated populations, as summarized by Tajima’s *D*, as a function of mutation rate (*μ*) and progeny skew (*Ψ*). For each parameter combination of *μ* and *Ψ* drawn from the prior distributions (10,000 in total), 1,000 replicates were simulated, with the mean given by the black dot and the standard deviation given by the gray bars. *Ψ* proportions are given on the X-axis, while the mean Tajima’s *D* is given on the Y-axis. Each panel corresponds to a different order of magnitude of mutation rate: (Α) 1e-6 to 9e-6, (B) 1e-7 to 9e-7, (C) 1e-8 to 9e-8, and (D) 1e-9 to 9e-9. As shown, the distribution of variation expected to be observed in the genome differs as a function of these two parameters - with a strong excess of rare mutations (negative *D*) observed as the WF model is initially violated at small values of *Ψ*, with a slight recovery towards 0 (and larger standard deviations) observed as *Ψ* gets larger, owing to the rapidly decreasing number of segregating sites associated with increased progeny skew.

**Figure S4.**
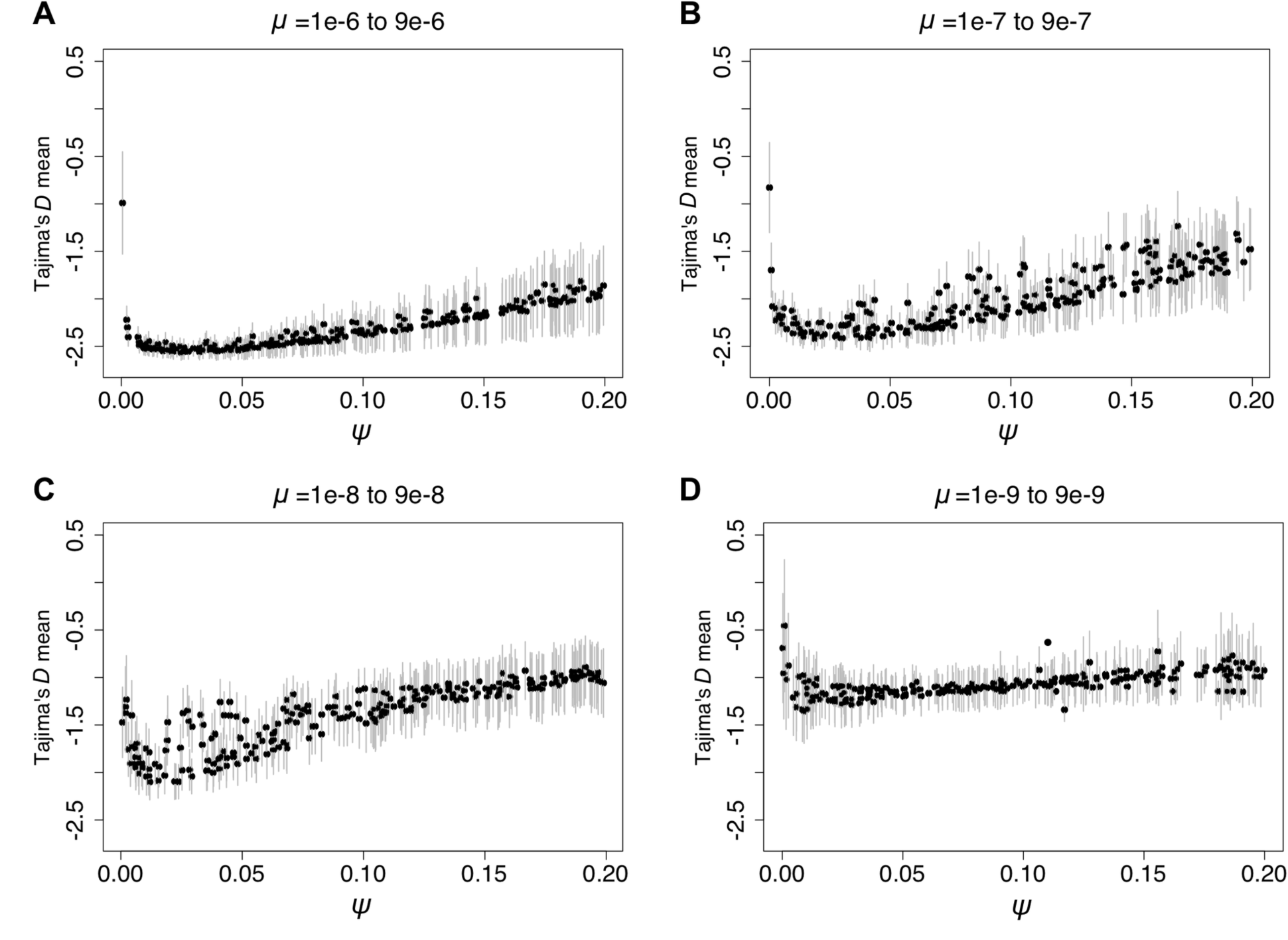
The shape of the site frequency spectrum (SFS), as summarized by Tajima’s *D,* in simulated populations after filtering out segregating sites under 2% frequency, as a function of mutation rate (*μ*) and progeny skew (*Ψ*). 1,000 parameters combinations were used. For each parameter combination of *μ* and *Ψ* drawn from the prior distributions, 1,000 replicates were simulated, with the mean given by the black dot and the standard deviation given by the gray bars. *Ψ* proportions are given on the X-axis, while the mean Tajima’s *D* is given on the Y-axis. Each panel corresponds to a different order of magnitude of mutation rate: (Α) 1e-6 to 9e-6, (B) 1e-7 to 9e-7, (C) 1e-8 to 9e-8, and (D) 1e-9 to 9e-9. As shown, the distribution of variation expected to be observed in the genome differs as a function of these two parameters - with a strong excess of rare mutations (negative *D*) observed as the WF model is initially violated at small values of *Ψ*, with a slight recovery towards 0 (and larger standard deviations) observed as *Ψ* gets larger, owing to the rapidly decreasing number of segregating sites associated with increased progeny skew.

**Figure S5.**
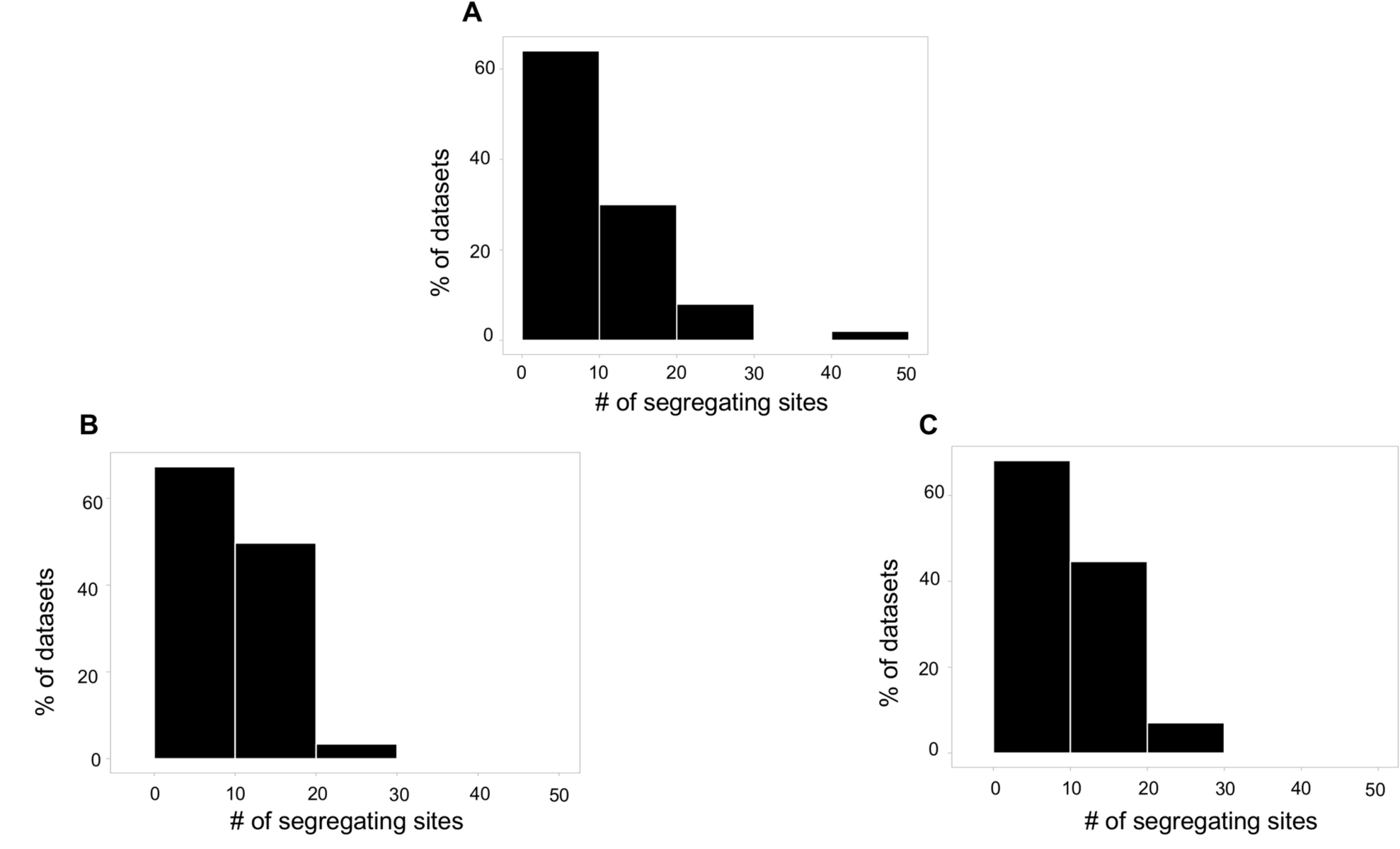
The distribution of segregating sites in real versus simulated datasets. Comparison of the number of segregating sites observed from (A) patient samples (Trauner *et al*. 2017), (B) simulated datasets with *μ* = 3.1e-8 and *Ψ* = 0.02, resulting in the same mean number of segregating sites as observed in patients, and (C) simulated datasets with *μ* = 3.7e-7 and *Ψ* = 0.18, also resulting in the same mean number of segregating sites as observed in patients. Thus, these two point estimates essentially demarcate the boundaries on the joint posterior ridge (Figure 3).

**Table S1.** Simulation results. Parameters simulated, ordered by descending mutation rate (*μ*) and corresponding values of progeny skew (*Ψ*) and bottleneck severity (*N2*). Given are the mean and standard deviation of the summary statistics from 1000 replicates.

| Parameters |  |  | Summary statistics |  |  |  |  |
| --- | --- | --- | --- | --- | --- | --- | --- |
| Mutation rate ($\mu$) | Progeny skew ($\psi$) | Bottle-neck ($N2$) | Tajima's $D$ mean | Tajima's $D$ sd | Seg. sites mean | Seg. Sites sd | % invariabl e genomes |
| 1.00323E-06 | 0.083741128 | 30 | -2.268823115 | 0.213071226 | 73.25 | 27.92044146 | 0 |
| 1.05258E-06 | 0.05424026 | 51 | -2.39378792 | 0.140975196 | 114.033 | 38.29559326 | 0 |
| 1.07427E-06 | 0.068391187 | 79 | -2.342687183 | 0.150033316 | 62.118 | 22.00859956 | 0 |
| 1.14948E-06 | 0.174968669 | 27 | -1.84867593 | 0.350954908 | 34.148 | 14.59140175 | 0 |
| 1.15893E-06 | 0.078976259 | 55 | -2.314091665 | 0.187904351 | 87.857 | 32.7311454 | 0 |
| 1.16072E-06 | 0.147004609 | 47 | -1.994103492 | 0.324323366 | 43.572 | 18.32666589 | 0 |
| 1.17846E-06 | 0.095707498 | 50 | -2.242540595 | 0.219702856 | 74.732 | 27.21637204 | 0 |
| 1.21809E-06 | 0.002234156 | 94 | -2.219989055 | 0.142920773 | 530.177 | 111.4160545 | 0 |
| 1.27435E-06 | 0.035230792 | 98 | -2.45132687 | 0.111998152 | 184.69 | 55.9886652 | 0 |
| 1.34490E-06 | 0.072224963 | 2 | -2.361174432 | 0.168335271 | 108.132 | 36.59953691 | 0 |
| 1.36631E-06 | 0.190031564 | 88 | -1.814642081 | 0.40537193 | 36.049 | 15.11837174 | 0 |
| 1.37186E-06 | 0.178587611 | 8 | -1.856448312 | 0.377166171 | 38.995 | 16.4323156 | 0 |
| 1.40763E-06 | 0.018259159 | 36 | -2.459363944 | 0.105067026 | 299.336 | 81.75943805 | 0 |
| 1.41886E-06 | 0.048682507 | 78 | -2.43334348 | 0.133550871 | 159.515 | 54.08471995 | 0 |
| 1.45413E-06 | 0.125929607 | 76 | -2.130629042 | 0.278288809 | 65.433 | 24.43773135 | 0 |
| 1.49860E-06 | 0.034752818 | 87 | -2.470499994 | 0.107467985 | 206.161 | 63.08506007 | 0 |
| 1.51763E-06 | 0.142547134 | 57 | -2.068036111 | 0.303344227 | 58.533 | 21.69316723 | 0 |
| 1.51902E-06 | 0.127470119 | 5 | -2.137500358 | 0.268938506 | 67.735 | 25.93629631 | 0 |
| 1.54224E-06 | 0.097141999 | 10 | -2.252101995 | 0.212272889 | 91.618 | 33.4446501 | 0 |
| 1.56157E-06 | 0.127268671 | 73 | -2.134927218 | 0.275793862 | 69.918 | 27.12203681 | 0 |
| 1.60310E-06 | 0.002535884 | 89 | -2.300181435 | 0.120396219 | 581.913 | 125.9045805 | 0 |
| 1.60552E-06 | 0.114838011 | 13 | -2.183291358 | 0.245635241 | 77.144 | 29.58409171 | 0 |
| 1.62498E-06 | 0.185445543 | 44 | -1.873943114 | 0.390190692 | 43.922 | 18.62054351 | 0 |
| 1.70401E-06 | 0.079764023 | 56 | -2.341980367 | 0.184901919 | 120.627 | 42.28086497 | 0 |
| 1.76944E-06 | 0.179382064 | 27 | -1.909361982 | 0.369374432 | 48.587 | 21.04122535 | 0 |
| 1.79775E-06 | 0.006932201 | 87 | -2.40224289 | 0.110270435 | 504.71 | 117.2538458 | 0 |
| 1.80560E-06 | 0.066056427 | 45 | -2.399866678 | 0.146367114 | 150.828 | 51.46002867 | 0 |
| 1.81127E-06 | 0.172076915 | 37 | -1.933554838 | 0.370411584 | 53.249 | 20.5200895 | 0 |
| 1.81152E-06 | 0.089215444 | 31 | -2.320938106 | 0.183317642 | 115.119 | 41.23101789 | 0 |
| 1.81640E-06 | 0.011295921 | 70 | -2.44570659 | 0.100980653 | 436.728 | 103.7460807 | 0 |
| 1.82390E-06 | 0.171978735 | 48 | -1.941706405 | 0.363939575 | 54.583 | 21.79017951 | 0 |
| 1.86518E-06 | 0.103034261 | 72 | -2.26005553 | 0.231646438 | 99.9 | 36.87109271 | 0 |
| 1.89217E-06 | 0.066288267 | 30 | -2.406201331 | 0.157309604 | 156.594 | 51.49237447 | 0 |
| 1.90714E-06 | 0.07621348 | 53 | -2.365034087 | 0.180909287 | 140.437 | 50.61689541 | 0 |
| 1.91685E-06 | 0.091231741 | 14 | -2.319495906 | 0.20669643 | 119.203 | 41.9865295 | 0 |
| 1.98106E-06 | 0.071256528 | 25 | -2.389881138 | 0.162727021 | 153.298 | 52.88578355 | 0 |
| 2.04999E-06 | 0.06206469 | 89 | -2.427399389 | 0.142401025 | 173.69 | 56.74013873 | 0 |
| 2.09647E-06 | 0.038639738 | 42 | -2.485880639 | 0.10741806 | 247.237 | 74.11912355 | 0 |
| 2.11204E-06 | 0.169388078 | 57 | -1.984453843 | 0.330080624 | 48.27233429 | 17.97727958 | 30.6 |
| 2.12986E-06 | 0.11706933 | 77 | -2.215467125 | 0.253808997 | 100.631 | 37.99700793 | 0 |
| 2.15406E-06 | 0.141870623 | 51 | -2.10605375 | 0.274222571 | 58.14 | 22.20408569 | 0 |
| 2.18101E-06 | 0.163422243 | 18 | -2.017817637 | 0.336968578 | 69.003 | 27.61317581 | 0 |
| 2.20091E-06 | 0.015172907 | 8 | -2.477645048 | 0.094911447 | 427.758 | 106.3263595 | 0 |
| 2.20482E-06 | 0.021126721 | 82 | -2.491883563 | 0.09687563 | 365.716 | 102.0283643 | 0 |
| 2.20725E-06 | 0.188675008 | 28 | -1.902430476 | 0.40017778 | 57.49 | 23.34714092 | 0 |
| 2.21559E-06 | 0.0871031 | 42 | -2.343760706 | 0.186671872 | 139.264 | 49.71911397 | 0 |
| 2.23573E-06 | 0.187063106 | 45 | -1.907741686 | 0.358805823 | 44.854 | 17.1103316 | 0 |
| 2.27509E-06 | 0.125403484 | 5 | -2.203743064 | 0.23813124 | 98.003 | 36.40753355 | 0 |
| 2.27948E-06 | 0.137483753 | 32 | -2.145546977 | 0.283584244 | 87.13 | 33.44340834 | 0 |
| 2.35400E-06 | 0.046762946 | 44 | -2.475339189 | 0.123211173 | 236.372 | 73.3149828 | 0 |
| 2.52429E-06 | 0.191032835 | 83 | -1.885597745 | 0.407044539 | 63.423 | 26.09231457 | 0 |
| 2.61389E-06 | 0.024278147 | 67 | -2.503128139 | 0.091279236 | 385.128 | 108.5069277 | 0 |
| 2.66469E-06 | 0.143581712 | 59 | -2.113962185 | 0.314452043 | 96.28 | 36.99459691 | 0 |
| 2.70701E-06 | 0.19942529 | 66 | -1.86045003 | 0.416596013 | 64.021 | 25.95500378 | 0 |
| 2.73336E-06 | 0.025876486 | 80 | -2.511972061 | 0.089679995 | 376.284 | 107.5730603 | 0 |
| 2.86631E-06 | 0.029798342 | 3 | -2.521767014 | 0.080400891 | 166.654 | 52.68561529 | 0 |
| 2.87173E-06 | 0.142357014 | 34 | -2.125391995 | 0.297032848 | 104.745 | 38.77612104 | 0 |
| 2.93144E-06 | 0.053739131 | 47 | -2.465139706 | 0.132375425 | 251.39 | 82.07500354 | 0 |
| 2.94178E-06 | 0.096296588 | 12 | -2.342917275 | 0.184995675 | 157.707 | 54.81443033 | 0 |
| 3.00818E-06 | 0.080858986 | 52 | -2.394854503 | 0.172760168 | 187.604 | 66.3320822 | 0 |
| 3.01815E-06 | 0.134974227 | 3 | -2.177646077 | 0.277538657 | 114.739 | 40.90511101 | 0 |
| 3.02499E-06 | 0.036913487 | 31 | -2.506608647 | 0.103410471 | 325.834 | 102.6217992 | 0 |
| 3.02691E-06 | 0.099126233 | 97 | -2.323489639 | 0.207079361 | 160.461 | 57.37567819 | 0 |
| 3.04722E-06 | 0.177489278 | 20 | -1.996511616 | 0.345804407 | 82.887 | 33.44183888 | 0 |
| 3.08878E-06 | 0.166632161 | 47 | -2.035520178 | 0.316529906 | 64.37974684 | 25.7113447 | 52.6 |
| 3.10911E-06 | 0.074397827 | 68 | -2.423287664 | 0.162287959 | 208.453 | 73.55282326 | 0 |
| 3.15283E-06 | 0.105716416 | 97 | -2.30220408 | 0.21672073 | 150.87 | 55.34531782 | 0 |
| 3.16156E-06 | 0.060129296 | 50 | -2.470586921 | 0.129254198 | 246.51 | 82.18757021 | 0 |
| 3.18711E-06 | 0.187231211 | 95 | -1.932400843 | 0.397556019 | 80.04 | 32.79457095 | 0 |
| 3.19924E-06 | 0.160557925 | 67 | -2.075862881 | 0.323439171 | 98.938 | 36.62460263 | 0 |
| 3.21256E-06 | 0.197806677 | 97 | -1.901179917 | 0.411388562 | 73.482 | 30.09819891 | 0 |
| 3.23557E-06 | 0.096667975 | 82 | -2.33429967 | 0.203058174 | 171.881 | 60.98852711 | 0 |
| 3.23876E-06 | 0.032477714 | 5 | -2.513252273 | 0.099558469 | 371.414 | 113.9971313 | 0 |
| 3.23960E-06 | 0.106816905 | 45 | -2.299007369 | 0.227105568 | 155.651 | 57.23310954 | 0 |
| 3.25363E-06 | 0.010197306 | 81 | -2.470389732 | 0.091743258 | 615.458 | 144.4888621 | 0 |
| 3.34148E-06 | 0.09036916 | 69 | -2.360354627 | 0.183471731 | 188.152 | 66.35187614 | 0 |
| 3.38046E-06 | 0.075316468 | 10 | -2.423214672 | 0.158800999 | 223.286 | 76.95137242 | 0 |
| 3.38234E-06 | 0.183560797 | 17 | -1.992111459 | 0.318425592 | 62.81481481 | 24.34369074 | 56.8 |
| 3.43922E-06 | 0.052317077 | 29 | -2.496508588 | 0.110215045 | 143.2914573 | 48.37521773 | 60.2 |
| 3.51100E-06 | 0.091190111 | 30 | -2.36341697 | 0.189148653 | 191.025 | 66.20401307 | 0 |
| 3.62777E-06 | 0.174054427 | 62 | -2.020730272 | 0.375769104 | 98.791 | 38.18973389 | 0 |
| 3.63591E-06 | 0.020922852 | 17 | -2.523624462 | 0.086263393 | 491.719 | 134.7929344 | 0 |
| 3.71752E-06 | 0.007424968 | 35 | -2.455211889 | 0.089901547 | 751.515 | 171.6138721 | 0 |
| 3.72700E-06 | 0.071519747 | 16 | -2.439368425 | 0.144560149 | 246.579 | 84.58926115 | 0 |
| 3.76115E-06 | 0.133389599 | 45 | -2.207183842 | 0.258449929 | 139.558 | 51.31061699 | 0 |
| 3.76891E-06 | 0.023000307 | 52 | -2.525610163 | 0.084925102 | 479.201 | 133.0190688 | 0 |
| 3.77969E-06 | 0.058721949 | 18 | -2.477337191 | 0.127971332 | 283.144 | 92.52348815 | 0 |
| 3.80658E-06 | 0.061662345 | 31 | -2.456041954 | 0.134864962 | 271.953 | 91.58006169 | 0 |
| 3.81465E-06 | 0.043844958 | 47 | -2.504175042 | 0.106799816 | 345.876 | 108.220702 | 0 |
| 3.85422E-06 | 0.045874048 | 89 | -2.509525385 | 0.095470736 | 162.163 | 51.36789943 | 0 |
| 3.89087E-06 | 0.157624492 | 50 | -2.100215392 | 0.317948113 | 117.258 | 45.17885085 | 0 |
| 3.92237E-06 | 0.105328979 | 66 | -2.325915846 | 0.216457163 | 183.751 | 65.4317225 | 0 |
| 3.93140E-06 | 0.097279879 | 22 | -2.348739672 | 0.199189136 | 197.806 | 70.31557896 | 0 |
| 4.05703E-06 | 0.081612311 | 27 | -2.401847125 | 0.167117424 | 231.15 | 77.99177018 | 0 |
| 4.06679E-06 | 0.052914714 | 99 | -2.49677306 | 0.114718297 | 320.979 | 100.8842713 | 0 |
| 4.08698E-06 | 0.162083695 | 39 | -2.082530637 | 0.338399342 | 119.814 | 46.42963617 | 0 |
| 4.11463E-06 | 0.147405055 | 97 | -2.161054228 | 0.286645225 | 136.814 | 51.00116197 | 0 |
| 4.13130E-06 | 0.071193318 | 91 | -2.450204823 | 0.139381082 | 261.729 | 88.2090494 | 0 |
| 4.13286E-06 | 0.178097374 | 72 | -2.005132374 | 0.379378644 | 104.374 | 41.08281784 | 0 |
| 4.13787E-06 | 0.183625187 | 25 | -1.98190994 | 0.385122469 | 105.096 | 42.00653917 | 0 |
| 4.18713E-06 | 0.115383656 | 91 | -2.279541427 | 0.233838275 | 177.233 | 61.31608576 | 0 |
| 4.21578E-06 | 0.063481085 | 74 | -2.465993109 | 0.131570546 | 283.843 | 93.59634944 | 0 |
| 4.22855E-06 | 0.044531836 | 24 | -2.514562421 | 0.10546776 | 365.052 | 116.1490934 | 0 |
| 4.26758E-06 | 0.064487967 | 61 | -2.466350062 | 0.137762942 | 292.162 | 96.33562326 | 0 |
| 4.27081E-06 | 0.057983672 | 56 | -2.480132515 | 0.126682888 | 310.77 | 98.58679656 | 0 |
| 4.27458E-06 | 0.051484956 | 70 | -2.502621768 | 0.119753276 | 341.951 | 109.9885725 | 0 |
| 4.28112E-06 | 0.081497163 | 85 | -2.414976591 | 0.172164667 | 245.994 | 82.79997464 | 0 |
| 4.33473E-06 | 0.18325412 | 16 | -1.996033612 | 0.346812486 | 73.077 | 28.78485085 | 0 |
| 4.34382E-06 | 0.089599967 | 85 | -2.389938479 | 0.160930254 | 126.775 | 44.33068446 | 0 |
| 4.35363E-06 | 0.031937187 | 16 | -2.533881587 | 0.088578631 | 441.809 | 130.8727863 | 0 |
| 4.38638E-06 | 0.170332639 | 1 | -2.043150239 | 0.357100011 | 121.465 | 47.98244565 | 0 |
| 4.40373E-06 | 0.180950653 | 97 | -2.02066542 | 0.34257131 | 112.818 | 44.79637922 | 0 |
| 4.44451E-06 | 0.009495684 | 39 | -2.48671751 | 0.089372565 | 763.837 | 181.69039 | 0 |
| 4.46435E-06 | 0.06388055 | 45 | -2.476451069 | 0.114443462 | 152.5642633 | 51.75499953 | 68.1 |
| 4.66398E-06 | 0.161534783 | 2 | -2.101108883 | 0.320353546 | 88.532 | 32.16074143 | 0 |
| 4.67889E-06 | 0.147050753 | 36 | -2.166447463 | 0.301217068 | 148.635 | 55.85450296 | 0 |
| 4.68696E-06 | 0.159150751 | 5 | -2.118221146 | 0.316422699 | 135.536 | 51.71722513 | 0 |
| 4.69022E-06 | 0.179421998 | 68 | -2.007102216 | 0.375179908 | 119.294 | 47.68985097 | 0 |
| 4.76545E-06 | 0.137827068 | 66 | -2.198064664 | 0.283620018 | 163.71 | 60.01127605 | 0 |
| 4.82616E-06 | 0.09864471 | 54 | -2.360936732 | 0.198095182 | 226.198 | 78.88992595 | 0 |
| 4.90490E-06 | 0.068348785 | 36 | -2.463883915 | 0.137071673 | 309.916 | 98.59704012 | 0 |
| 4.92672E-06 | 0.037491836 | 45 | -2.53464184 | 0.089585124 | 437.73 | 134.1963254 | 0 |
| 4.96090E-06 | 0.194931247 | 66 | -1.943824358 | 0.406728845 | 111.012 | 45.51902721 | 0 |
| 4.99886E-06 | 0.084205461 | 81 | -2.415349811 | 0.164869693 | 271.626 | 90.2507802 | 0 |
| 5.03965E-06 | 0.010818434 | 41 | -2.502006876 | 0.084952354 | 765.118 | 187.5172814 | 0 |
| 5.08458E-06 | 0.092243176 | 67 | -2.395655224 | 0.168627216 | 137.778 | 50.38058077 | 0 |
| 5.19177E-06 | 0.081319929 | 51 | -2.423738507 | 0.158052048 | 279.023 | 94.15976068 | 0 |
| 5.19312E-06 | 0.177636988 | 13 | -2.014820856 | 0.373052196 | 131.709 | 50.46064403 | 0 |
| 5.28653E-06 | 0.159011493 | 5 | -2.111142553 | 0.312880421 | 151.001 | 57.19405703 | 0 |
| 5.33565E-06 | 0.036433081 | 70 | -2.536428429 | 0.093074865 | 466.303 | 140.9724811 | 0 |
| 5.34859E-06 | 0.032126559 | 79 | -2.536982301 | 0.092139694 | 502.513 | 146.1721866 | 0 |
| 5.34876E-06 | 0.074147757 | 73 | -2.446396385 | 0.141400751 | 307.527 | 98.73268666 | 0 |
| 5.37165E-06 | 0.070847357 | 98 | -2.464197299 | 0.14237841 | 318.068 | 106.417687 | 0 |
| 5.39080E-06 | 0.148057423 | 61 | -2.17056632 | 0.291890977 | 164.095 | 58.76678918 | 0 |
| 5.42598E-06 | 0.033790744 | 76 | -2.537953881 | 0.090863628 | 490.955 | 145.8321981 | 0 |
| 5.45932E-06 | 0.150455894 | 94 | -2.138987203 | 0.303190167 | 163.692 | 61.6935753 | 0 |
| 5.49750E-06 | 0.013493694 | 73 | -2.51213801 | 0.084049365 | 745.896 | 194.3784818 | 0 |
| 5.50896E-06 | 0.148013485 | 41 | -2.179766751 | 0.27981175 | 171.601 | 64.51595235 | 0 |
| 5.52108E-06 | 0.017269882 | 8 | -2.530306408 | 0.085229634 | 680.684 | 183.9934285 | 0 |
| 5.59879E-06 | 0.079697476 | 68 | -2.42298135 | 0.161658037 | 302.269 | 106.0762877 | 0 |
| 5.61346E-06 | 0.039363853 | 66 | -2.533772282 | 0.099950806 | 457.639 | 145.3699457 | 0 |
| 5.62852E-06 | 0.134515261 | 5 | -2.229571912 | 0.254274997 | 187.168 | 68.44351167 | 0 |
| 5.63844E-06 | 0.165350342 | 49 | -2.092404381 | 0.342592855 | 154.336 | 57.47631365 | 0 |
| 5.64346E-06 | 0.197158598 | 52 | -1.948358035 | 0.406635491 | 120.541 | 47.30573064 | 0 |
| 5.66941E-06 | 0.014846757 | 37 | -2.527392022 | 0.084809703 | 731.887 | 186.8805516 | 0 |
| 5.77125E-06 | 0.039009255 | 20 | -2.534196812 | 0.096215357 | 475.949095 | 151.0681737 | 11.6 |
| 5.79828E-06 | 0.102032778 | 31 | -2.347687999 | 0.215802247 | 252.055 | 87.4202853 | 0 |
| 5.81859E-06 | 0.018901769 | 41 | -2.536851337 | 0.081223929 | 670.963 | 187.6406316 | 0 |
| 5.86554E-06 | 0.119172716 | 42 | -2.289135668 | 0.247164645 | 224.895 | 80.54838308 | 0 |
| 5.87513E-06 | 0.009945494 | 12 | -2.529829888 | 0.065175172 | 354.518 | 90.49086292 | 0 |
| 5.91748E-06 | 0.108639825 | 8 | -2.330618799 | 0.21801644 | 245.334 | 89.73793702 | 0 |
| 5.95191E-06 | 0.163364246 | 88 | -2.0916936 | 0.343509769 | 159.667 | 63.38283281 | 0 |
| 5.96091E-06 | 0.087444079 | 18 | -2.41819886 | 0.161575551 | 153.277 | 55.05082074 | 0 |
| 5.99521E-06 | 0.181327866 | 30 | -2.035142738 | 0.356564966 | 144.429 | 55.8730414 | 0 |
| 6.01869E-06 | 0.177907004 | 55 | -2.06257483 | 0.326697733 | 147.444 | 57.72470013 | 0 |
| 6.05063E-06 | 0.002877447 | 32 | -2.401946034 | 0.096067514 | 1198.869 | 255.7148841 | 0 |
| 6.07731E-06 | 0.01955547 | 1 | -2.541352169 | 0.080011363 | 681.491 | 179.4129955 | 0 |
| 6.14328E-06 | 0.060748557 | 55 | -2.49084606 | 0.129369442 | 383.209 | 129.8532063 | 0 |
| 6.21723E-06 | 0.043054603 | 2 | -2.532038626 | 0.097144385 | 474.703 | 147.9193257 | 0 |
| 6.22292E-06 | 0.051537806 | 68 | -2.522713259 | 0.107135081 | 422.113 | 136.9644972 | 0 |
| 6.28351E-06 | 0.04833965 | 58 | -2.528331181 | 0.102515432 | 449.096 | 145.2344237 | 0 |
| 6.28653E-06 | 0.081165105 | 44 | -2.431136744 | 0.155267755 | 321.523 | 106.133132 | 0 |
| 6.32747E-06 | 0.193498773 | 51 | -2.01025924 | 0.377904736 | 138.227 | 54.61023484 | 0 |
| 6.34468E-06 | 0.027728583 | 71 | -2.553733432 | 0.080758412 | 581.877 | 172.5323601 | 0 |
| 6.34891E-06 | 0.09256866 | 32 | -2.398773229 | 0.179032918 | 295.067 | 100.3194124 | 0 |
| 6.36937E-06 | 0.142826501 | 50 | -2.200359147 | 0.272622369 | 120.961 | 46.28833643 | 0 |
| 6.39225E-06 | 0.000498444 | 97 | -0.990254509 | 0.538172921 | 4711.724 | 1154.140446 | 0 |
| 6.42691E-06 | 0.049874555 | 75 | -2.520309606 | 0.10979855 | 442.701 | 142.79228 | 0 |
| 6.42861E-06 | 0.063886961 | 7 | -2.490548155 | 0.131109928 | 390.055 | 130.0545045 | 0 |
| 6.45845E-06 | 0.039415139 | 19 | -2.541081406 | 0.094079965 | 511.883 | 159.5622745 | 0 |
| 6.50933E-06 | 0.061705597 | 79 | -2.493064687 | 0.123437591 | 389.011 | 123.4561878 | 0 |
| 6.52081E-06 | 0.179715526 | 95 | -2.052669741 | 0.347732189 | 158.22 | 61.28929742 | 0 |
| 6.61941E-06 | 0.013406386 | 49 | -2.527695803 | 0.076238044 | 840.509 | 207.1711655 | 0 |
| 6.67778E-06 | 0.025203094 | 41 | -2.552025842 | 0.083223018 | 643.11 | 184.4100349 | 0 |
| 6.71129E-06 | 0.184675108 | 81 | -2.033468914 | 0.36471217 | 157.084 | 59.45143747 | 0 |
| 6.74110E-06 | 0.012953563 | 72 | -2.529307254 | 0.078355505 | 870.143 | 224.0870893 | 0 |
| 6.77895E-06 | 0.042308741 | 80 | -2.541125362 | 0.090781076 | 497.057 | 147.9309884 | 0 |
| 6.78473E-06 | 0.180034226 | 47 | -2.037501557 | 0.354121272 | 163.303 | 62.91191541 | 0 |
| 6.80869E-06 | 0.146559555 | 21 | -2.19406396 | 0.284609993 | 122.693 | 45.58109682 | 0 |
| 6.82733E-06 | 0.056909954 | 59 | -2.513305204 | 0.113574939 | 422.41 | 134.0480988 | 0 |
| 6.83593E-06 | 0.076868611 | 49 | -2.442675613 | 0.160653802 | 350.257 | 118.3542592 | 0 |
| 6.84453E-06 | 0.025937913 | 15 | -2.556708238 | 0.078860856 | 644.687 | 193.0205065 | 0 |
| 6.86818E-06 | 0.088019082 | 47 | -2.420797254 | 0.179780326 | 320.616 | 104.4551072 | 0 |
| 6.90801E-06 | 0.007457312 | 75 | -2.48893408 | 0.080148005 | 1048.332 | 239.3597562 | 0 |
| 6.95230E-06 | 0.186057268 | 78 | -2.018813781 | 0.350502131 | 157.646 | 60.18999518 | 0 |
| 6.98195E-06 | 0.150790823 | 8 | -2.173780372 | 0.296051761 | 201.855 | 76.85593542 | 0 |
| 7.01129E-06 | 0.117400419 | 99 | -2.317364359 | 0.223678581 | 260.326 | 92.96620052 | 0 |
| 7.02328E-06 | 0.062670765 | 68 | -2.495649631 | 0.12599538 | 408.589 | 129.633317 | 0 |
| 7.03960E-06 | 0.076898129 | 49 | -2.458608975 | 0.144983134 | 369.077 | 124.3281326 | 0 |
| 7.04110E-06 | 0.02039321 | 27 | -2.551816517 | 0.073694819 | 739.155 | 201.868308 | 0 |
| 7.07798E-06 | 0.09952533 | 11 | -2.367716285 | 0.202708985 | 298.488 | 106.8031028 | 0 |
| 7.16578E-06 | 0.164836135 | 34 | -2.112573648 | 0.324466583 | 189.619 | 73.83079863 | 0 |
| 7.16907E-06 | 0.178343668 | 30 | -2.06332979 | 0.329153933 | 112.1597938 | 39.12594559 | 80.6 |
| 7.17772E-06 | 0.138474662 | 21 | -2.229724935 | 0.26582104 | 230.211 | 79.97094367 | 0 |
| 7.18952E-06 | 0.051417423 | 58 | -2.529305237 | 0.107858519 | 474.913 | 150.5340624 | 0 |
| 7.21034E-06 | 0.131057148 | 41 | -2.263690883 | 0.236809421 | 238.425 | 84.12810418 | 0 |
| 7.21766E-06 | 0.150426253 | 89 | -2.183733977 | 0.30257552 | 210.568 | 80.25684106 | 0 |
| 7.22276E-06 | 0.065801281 | 27 | -2.487010759 | 0.135017996 | 395.223 | 131.0033326 | 0 |
| 7.28827E-06 | 0.054523495 | 28 | -2.512510853 | 0.117947502 | 447.441 | 148.2902712 | 0 |
| 7.34886E-06 | 0.048310702 | 34 | -2.530371191 | 0.106199965 | 494.59 | 158.0397053 | 0 |
| 7.36520E-06 | 0.051301492 | 95 | -2.51914289 | 0.113994345 | 472.227 | 147.7385398 | 0 |
| 7.39311E-06 | 0.048410189 | 44 | -2.543515174 | 0.089002604 | 222.213 | 75.67403908 | 0 |
| 7.39443E-06 | 0.142488828 | 65 | -2.223760785 | 0.272916523 | 228.38 | 82.59871759 | 0 |
| 7.41247E-06 | 0.082466592 | 26 | -2.445896823 | 0.155609023 | 350.58 | 127.4844614 | 0 |
| 7.42439E-06 | 0.138494135 | 4 | -2.226955385 | 0.279720187 | 232.196 | 84.05707094 | 0 |
| 7.50443E-06 | 0.074608316 | 53 | -2.467081169 | 0.144974333 | 380.4653266 | 126.8435385 | 0.5 |
| 7.51767E-06 | 0.137078511 | 72 | -2.245032333 | 0.257150794 | 236.178 | 86.07827708 | 0 |
| 7.59875E-06 | 0.04001344 | 92 | -2.548634304 | 0.094289806 | 549.201 | 174.9091301 | 0 |
| 7.60124E-06 | 0.197655797 | 56 | -1.971468748 | 0.394631373 | 160.586 | 61.63470971 | 0 |
| 7.64061E-06 | 0.075298103 | 79 | -2.459775663 | 0.156429417 | 386.462 | 132.3871252 | 0 |
| 7.68735E-06 | 0.086217638 | 71 | -2.447923552 | 0.143747413 | 179.422 | 63.82486608 | 0 |
| 7.73333E-06 | 0.079299325 | 79 | -2.452550733 | 0.15307327 | 376.58 | 127.8387983 | 0 |
| 7.77212E-06 | 0.196554711 | 63 | -1.984723902 | 0.392655194 | 164.695 | 63.20779382 | 0 |
| 7.80427E-06 | 0.011199664 | 63 | -2.520420468 | 0.077848821 | 966.524 | 235.7493355 | 0 |
| 7.86550E-06 | 0.179721627 | 71 | -2.06962471 | 0.341131207 | 181.521 | 69.42190548 | 0 |
| 7.88451E-06 | 0.080996722 | 35 | -2.449756283 | 0.153657455 | 383.829 | 133.0214662 | 0 |
| 7.88638E-06 | 0.1666448 | 59 | -2.120404744 | 0.322095976 | 201.396 | 75.09448959 | 0 |
| 7.92294E-06 | 0.067269241 | 93 | -2.485810753 | 0.13481249 | 418.795 | 138.7381103 | 0 |
| 7.98441E-06 | 0.061223256 | 13 | -2.506154325 | 0.119019726 | 451.1435794 | 146.2317971 | 1.1 |
| 7.99095E-06 | 0.089447718 | 53 | -2.440670425 | 0.149941216 | 179.837 | 65.68188899 | 0 |
| 7.99285E-06 | 0.050713501 | 97 | -2.534753003 | 0.106458715 | 495.403 | 155.9264623 | 0 |
| 7.99901E-06 | 0.050967349 | 89 | -2.528929069 | 0.103038136 | 497.634 | 154.4307252 | 0 |
| 8.00013E-06 | 0.128142071 | 83 | -2.277511989 | 0.24697584 | 268.947 | 96.30920857 | 0 |
| 8.00249E-06 | 0.11790393 | 55 | -2.322167818 | 0.227962689 | 288.982 | 97.69893939 | 0 |
| 8.00836E-06 | 0.148077971 | 87 | -2.199919025 | 0.284032144 | 136.619 | 48.85972894 | 0 |
| 8.11586E-06 | 0.167457995 | 87 | -2.132365478 | 0.320822963 | 124.238 | 46.64501166 | 0 |
| 8.13033E-06 | 0.021430876 | 75 | -2.551283793 | 0.079865134 | 773.333 | 207.6434538 | 0 |
| 8.17407E-06 | 0.125348653 | 80 | -2.284101759 | 0.239945517 | 276.9866518 | 96.61832147 | 10.1 |
| 8.27083E-06 | 0.074989706 | 45 | -2.464181313 | 0.152911371 | 413.274 | 143.7678619 | 0 |
| 8.29799E-06 | 0.024720669 | 88 | -2.566448744 | 0.079759978 | 719.667 | 211.7515214 | 0 |
| 8.30624E-06 | 0.104965581 | 65 | -2.363483165 | 0.200270093 | 321.74 | 112.5569521 | 0 |
| 8.44438E-06 | 0.186754651 | 80 | -2.018784115 | 0.383265374 | 188.768 | 72.4658366 | 0 |
| 8.46443E-06 | 0.101071854 | 5 | -2.378379641 | 0.2010005 | 335.763 | 113.2412664 | 0 |
| 8.46703E-06 | 0.165195627 | 75 | -2.129612837 | 0.31867126 | 216.681 | 83.37253876 | 0 |
| 8.46893E-06 | 0.032363037 | 37 | -2.55913469 | 0.082819055 | 653.552 | 201.7541525 | 0 |
| 8.49289E-06 | 0.052155964 | 70 | -2.529301194 | 0.10687454 | 511.329 | 168.0613977 | 0 |
| 8.49640E-06 | 0.011483327 | 93 | -2.524171765 | 0.077585068 | 995.999 | 253.6725029 | 0 |
| 8.50168E-06 | 0.157491798 | 58 | -2.150882271 | 0.3135751 | 224.714 | 82.46358255 | 0 |
| 8.59353E-06 | 0.119391729 | 2 | -2.310786869 | 0.233170398 | 301.436 | 110.8122613 | 0 |
| 8.63529E-06 | 0.112601893 | 34 | -2.331966464 | 0.228850405 | 313.448 | 111.5213805 | 0 |
| 8.64280E-06 | 0.128128245 | 19 | -2.258019263 | 0.255049042 | 281.851 | 100.7786641 | 0 |
| 8.72579E-06 | 0.133745637 | 41 | -2.247429925 | 0.277316686 | 270.206 | 100.7595854 | 0 |
| 8.73167E-06 | 0.021543106 | 22 | -2.552923449 | 0.082063861 | 785.933 | 223.020202 | 0 |
| 8.74277E-06 | 0.017176358 | 80 | -2.549018151 | 0.076780648 | 864.539 | 242.7295081 | 0 |
| 8.77671E-06 | 0.116429936 | 1 | -2.323459329 | 0.237454452 | 310.695 | 109.1745715 | 0 |
| 8.79343E-06 | 0.103772443 | 78 | -2.376890594 | 0.188639046 | 335.424 | 119.675923 | 0 |
| 8.80503E-06 | 0.009299307 | 97 | -2.515949453 | 0.075308786 | 1085.272 | 272.3681046 | 0 |
| 8.81455E-06 | 0.039320445 | 79 | -2.555134061 | 0.092649516 | 612.422 | 203.4150181 | 0 |
| 8.84282E-06 | 0.113147577 | 88 | -2.342495118 | 0.20960697 | 319.741 | 108.2580053 | 0 |
| 8.89345E-06 | 0.045187261 | 8 | -2.543956231 | 0.098464532 | 560.6888658 | 173.7109308 | 3.9 |
| 1.01007E-07 | 0.037207555 | 42 | -2.051728712 | 0.197389059 | 21.65 | 8.301229452 | 0 |
| 1.06501E-07 | 0.043483318 | 60 | -2.012139803 | 0.212294591 | 19.3993994 | 7.609036372 | 0.1 |
| 1.11486E-07 | 0.038947139 | 4 | -2.059089214 | 0.203824726 | 22.616 | 8.495217049 | 0 |
| 1.16237E-07 | 0.086858593 | 96 | -1.690919447 | 0.28940054 | 9.419715447 | 4.341671266 | 1.6 |
| 1.30883E-07 | 0.082461751 | 26 | -1.76546852 | 0.281696443 | 11.30251256 | 5.274167687 | 0.5 |
| 1.31513E-07 | 0.169136567 | 32 | -1.235420133 | 0.365639957 | 4.790486976 | 2.412702611 | 11.7 |
| 1.31540E-07 | 0.073409472 | 70 | -1.865148747 | 0.25371201 | 13.37951807 | 5.777146291 | 0.4 |
| 1.31617E-07 | 0.009450759 | 77 | -2.125155336 | 0.185832109 | 76.434 | 19.25191628 | 0 |
| 1.34829E-07 | 0.000769518 | 77 | -1.697173636 | 0.284645295 | 136.115 | 30.2327934 | 0 |
| 1.38570E-07 | 0.084078144 | 44 | -1.780101152 | 0.272122944 | 11.87638191 | 5.493346531 | 0.5 |
| 1.43412E-07 | 0.105046852 | 85 | -1.637557995 | 0.301087978 | 9.364646465 | 4.619753891 | 1 |
| 1.46392E-07 | 0.040145492 | 17 | -2.135284109 | 0.184647437 | 28.418 | 10.39332946 | 0 |
| 1.48040E-07 | 0.005775423 | 78 | -2.098596342 | 0.182797238 | 98.163 | 24.08057942 | 0 |
| 1.51325E-07 | 0.105622891 | 79 | -1.670288298 | 0.312594453 | 9.773373984 | 4.550212945 | 1.6 |
| 1.53729E-07 | 0.034102996 | 71 | -2.178654623 | 0.172540664 | 34.491 | 11.81259736 | 0 |
| 1.55198E-07 | 0.032455678 | 97 | -2.191126512 | 0.173775225 | 37.506 | 12.53375678 | 0 |
| 1.56929E-07 | 0.057293669 | 13 | -2.042616833 | 0.20408599 | 20.968 | 8.100283365 | 0 |
| 1.58551E-07 | 0.090848716 | 65 | -1.769475169 | 0.281742653 | 12.20523139 | 5.734485422 | 0.6 |
| 1.59398E-07 | 0.140382942 | 9 | -1.456672525 | 0.367980397 | 7.033057851 | 3.521384245 | 3.2 |
| 1.61103E-07 | 0.070249371 | 69 | -1.936668307 | 0.248297836 | 17.00801603 | 7.294076125 | 0.2 |
| 1.61814E-07 | 0.147711891 | 83 | -1.431511401 | 0.330481489 | 6.719334719 | 3.289408534 | 3.8 |
| 1.64562E-07 | 0.034303216 | 59 | -2.204674548 | 0.164424981 | 37.408 | 12.50815694 | 0 |
| 1.64607E-07 | 0.04166588 | 29 | -2.148898671 | 0.186285207 | 30.243 | 10.84147077 | 0 |
| 1.67115E-07 | 0.042028676 | 83 | -2.158735481 | 0.171895398 | 30.332 | 11.00776757 | 0 |
| 1.72210E-07 | 0.160113083 | 69 | -1.396790625 | 0.351619853 | 6.412147505 | 3.224058574 | 7.8 |
| 1.75415E-07 | 0.104000951 | 12 | -1.740389559 | 0.287249414 | 11.67507568 | 5.48417138 | 0.9 |
| 1.81932E-07 | 0.14681816 | 88 | -1.461909975 | 0.35936019 | 7.61761658 | 3.760172995 | 3.5 |
| 1.83192E-07 | 0.021043343 | 23 | -2.251400713 | 0.149714884 | 60.754 | 17.8018648 | 0 |
| 1.88713E-07 | 0.156515898 | 48 | -1.392182677 | 0.381506499 | 7.013569937 | 3.69342713 | 4.2 |
| 1.90009E-07 | 0.021541516 | 89 | -2.248712907 | 0.155683169 | 59.158 | 17.47599629 | 0 |
| 1.91987E-07 | 0.010356717 | 30 | -2.202204033 | 0.168264635 | 97.794 | 24.93181019 | 0 |
| 1.93567E-07 | 0.020632953 | 51 | -2.252492364 | 0.155272094 | 63.037 | 18.30378812 | 0 |
| 2.03309E-07 | 0.157051872 | 34 | -1.456888632 | 0.381325772 | 7.923879041 | 4.091776361 | 4.1 |
| 2.18721E-07 | 0.086025658 | 74 | -1.924202682 | 0.263315314 | 17.60760761 | 7.361151764 | 0.1 |
| 2.19485E-07 | 0.032888251 | 27 | -2.256192839 | 0.153005715 | 49.268 | 15.75983974 | 0 |
| 2.19679E-07 | 0.154142694 | 77 | -1.495808435 | 0.371576186 | 8.616803279 | 4.153797697 | 2.4 |
| 2.22485E-07 | 0.126848726 | 43 | -1.647251953 | 0.32377229 | 10.78947368 | 5.211262308 | 1.2 |
| 2.25032E-07 | 0.155844325 | 48 | -1.480786984 | 0.373873978 | 8.399795501 | 4.352986946 | 2.2 |
| 2.29831E-07 | 0.030077451 | 71 | -2.27335215 | 0.143059314 | 55.338 | 18.12904902 | 0 |
| 2.31006E-07 | 0.016107215 | 84 | -2.263564921 | 0.153668533 | 84.815 | 22.90955309 | 0 |
| 2.42615E-07 | 0.094599317 | 40 | -1.899536017 | 0.269312799 | 17.18336673 | 7.410681378 | 0.2 |
| 2.44791E-07 | 0.024944146 | 43 | -2.286677176 | 0.146057109 | 68.256 | 21.01609241 | 0 |
| 2.45494E-07 | 0.029922845 | 11 | -2.284437618 | 0.152107372 | 58.574 | 18.39992068 | 0 |
| 2.48791E-07 | 0.157492765 | 73 | -1.527458089 | 0.351890945 | 9.229441624 | 4.561475544 | 1.5 |
| 2.52080E-07 | 0.193546788 | 21 | -1.31359069 | 0.38831406 | 7.111461619 | 3.590365041 | 4.9 |
| 2.54351E-07 | 0.032071962 | 69 | -2.281012588 | 0.151163693 | 56.094 | 18.23056547 | 0 |
| 2.55051E-07 | 0.115715145 | 62 | -1.808019754 | 0.283378053 | 14.54929577 | 6.527043788 | 0.6 |
| 2.56097E-07 | 0.068202216 | 46 | -2.099016583 | 0.221928491 | 27.858 | 11.27656849 | 0 |
| 2.57110E-07 | 0.130054995 | 61 | -1.693241351 | 0.337814069 | 12.3919598 | 5.889930677 | 0.5 |
| 2.60855E-07 | 0.134972889 | 90 | -1.676219507 | 0.334868692 | 11.80524723 | 5.582205921 | 0.9 |
| 2.65494E-07 | 0.159940203 | 68 | -1.522496241 | 0.359625464 | 9.347560976 | 4.465207894 | 1.6 |
| 2.66887E-07 | 0.137423897 | 27 | -1.656033391 | 0.335390019 | 11.62273642 | 5.640231523 | 0.6 |
| 2.67293E-07 | 0.178332191 | 26 | -1.429277855 | 0.390284363 | 8.269270298 | 4.152624633 | 2.7 |
| 2.67657E-07 | 0.01171703 | 22 | -2.26184749 | 0.151632922 | 115.415 | 29.1269833 | 0 |
| 2.77101E-07 | 0.089815698 | 84 | -1.97863907 | 0.262622641 | 21.281 | 8.637629747 | 0 |
| 2.84542E-07 | 0.159657855 | 67 | -1.551669886 | 0.360559259 | 10.24696356 | 4.913614534 | 1.2 |
| 2.86705E-07 | 0.076434857 | 35 | -2.080676078 | 0.217220878 | 27.147 | 10.96565715 | 0 |
| 2.95952E-07 | 0.009966019 | 61 | -2.262246359 | 0.151047832 | 135.232 | 32.51974338 | 0 |
| 2.98885E-07 | 0.121334637 | 44 | -1.800200899 | 0.322363033 | 15.65295888 | 6.952934414 | 0.3 |
| 3.00067E-07 | 0.194122681 | 2 | -1.381106528 | 0.400836709 | 8.141675284 | 4.106449818 | 3.3 |
| 3.02974E-07 | 0.089504202 | 68 | -2.002698716 | 0.24189679 | 23.6 | 9.763879011 | 0 |
| 3.20559E-07 | 0.088571362 | 72 | -2.015069109 | 0.256241002 | 24.705 | 10.17799423 | 0 |
| 3.22863E-07 | 0.097689633 | 25 | -1.993466538 | 0.243670378 | 22.576 | 9.420348559 | 0 |
| 3.26118E-07 | 0.036476882 | 14 | -2.303960911 | 0.146584594 | 62.736 | 21.17547413 | 0 |
| 3.27505E-07 | 0.005282555 | 41 | -2.179380302 | 0.166583287 | 186.832 | 40.77330301 | 0 |
| 3.29323E-07 | 0.050551651 | 60 | -2.253900919 | 0.177992937 | 46.379 | 16.37392636 | 0 |
| 3.29820E-07 | 0.067238279 | 28 | -2.163622197 | 0.191617613 | 34.587 | 12.93598743 | 0 |
| 3.30007E-07 | 0.156793492 | 89 | -1.617907817 | 0.357900941 | 11.94869215 | 5.81767139 | 0.6 |
| 3.35742E-07 | 0.167702541 | 54 | -1.548066791 | 0.381756124 | 10.81010101 | 5.146027259 | 1 |
| 3.41295E-07 | 0.173539696 | 52 | -1.533585629 | 0.380551779 | 10.71890799 | 5.294739619 | 1.1 |
| 3.42671E-07 | 0.082308406 | 83 | -2.089385281 | 0.218527615 | 28.736 | 11.30673013 | 0 |
| 3.42882E-07 | 0.07431508 | 8 | -2.121981096 | 0.225706387 | 32.72 | 12.81154697 | 0 |
| 3.45360E-07 | 0.042748883 | 49 | -2.288643438 | 0.158097015 | 56.221 | 19.0995013 | 0 |
| 3.58661E-07 | 0.07559624 | 28 | -2.134153711 | 0.208705032 | 33.103 | 13.04954718 | 0 |
| 3.59921E-07 | 0.125055574 | 99 | -1.829067947 | 0.310160581 | 17.93881645 | 7.852765852 | 0.3 |
| 3.61200E-07 | 0.166728483 | 96 | -1.59625845 | 0.370273383 | 12.28830645 | 5.890828558 | 0.8 |
| 3.61990E-07 | 0.115642209 | 19 | -1.894221514 | 0.304138975 | 19.73073073 | 8.826686091 | 0.1 |
| 3.67849E-07 | 0.088095933 | 67 | -2.072838478 | 0.234517588 | 28.558 | 10.95968753 | 0 |
| 3.71272E-07 | 0.180958839 | 47 | -1.509631867 | 0.405640517 | 11.18705763 | 5.698781241 | 1.1 |
| 3.71916E-07 | 0.185370158 | 53 | -1.470742075 | 0.395973823 | 10.56275304 | 5.375005885 | 1.2 |
| 3.75708E-07 | 0.059146015 | 60 | -2.242430884 | 0.17534194 | 45.729 | 16.15718458 | 0 |
| 3.79205E-07 | 0.126706296 | 27 | -1.841558912 | 0.298893032 | 18.514 | 7.936043396 | 0 |
| 3.81526E-07 | 0.012531531 | 55 | -2.309963075 | 0.145753473 | 143.198 | 36.64332858 | 0 |
| 3.81531E-07 | 0.184663853 | 87 | -1.482191711 | 0.425535539 | 10.89292929 | 5.422181394 | 1 |
| 3.84650E-07 | 0.035590645 | 80 | -2.327969953 | 0.151926452 | 72.516 | 22.81864226 | 0 |
| 3.85416E-07 | 0.175953718 | 53 | -1.563247984 | 0.378332212 | 11.71039354 | 5.718645999 | 0.9 |
| 3.86830E-07 | 0.106141973 | 46 | -1.974307014 | 0.262384794 | 23.65 | 9.968092238 | 0 |
| 3.88749E-07 | 0.079953214 | 58 | -2.147676944 | 0.205784745 | 33.827 | 13.18950469 | 0 |
| 3.88788E-07 | 0.04436575 | 64 | -2.310322328 | 0.154710767 | 61.693 | 20.85727156 | 0 |
| 3.90501E-07 | 0.058404112 | 96 | -2.240043769 | 0.172156923 | 47.056 | 17.70271872 | 0 |
| 3.91166E-07 | 0.138171295 | 90 | -1.790899531 | 0.327033394 | 17.18336673 | 7.311759663 | 0.2 |
| 3.92821E-07 | 0.003985377 | 82 | -2.168462609 | 0.162018642 | 224.207 | 47.82461024 | 0 |
| 3.93391E-07 | 0.041200387 | 66 | -2.312058383 | 0.156169676 | 66.356 | 21.61935847 | 0 |
| 4.02929E-07 | 0.169936849 | 3 | -1.603446846 | 0.381425273 | 13.05236657 | 6.586679873 | 0.7 |
| 4.11887E-07 | 0.06560143 | 87 | -2.215498124 | 0.188134209 | 43.547 | 16.05270944 | 0 |
| 4.14708E-07 | 0.110029495 | 25 | -1.959219285 | 0.266771854 | 23.897 | 10.10467691 | 0 |
| 4.18398E-07 | 0.171023922 | 9 | -1.628216432 | 0.35684027 | 13.5427996 | 6.409924184 | 0.7 |
| 4.20551E-07 | 0.090051366 | 33 | -2.096146008 | 0.224228824 | 31.441 | 12.4585607 | 0 |
| 4.25340E-07 | 0.137870962 | 64 | -1.812475466 | 0.31592473 | 18.34068136 | 8.114536675 | 0.2 |
| 4.26358E-07 | 0.166239545 | 10 | -1.662961204 | 0.374864082 | 14.6252505 | 6.943376348 | 0.2 |
| 4.28101E-07 | 0.158786361 | 54 | -1.70345215 | 0.351579083 | 15.60621242 | 7.121825798 | 0.2 |
| 4.30122E-07 | 0.141437011 | 3 | -1.781114828 | 0.325992818 | 17.60140562 | 7.776058995 | 0.4 |
| 4.34453E-07 | 0.18849351 | 70 | -1.524455338 | 0.407446459 | 11.95766129 | 5.741155772 | 0.8 |
| 4.42454E-07 | 0.017329756 | 89 | -2.347727504 | 0.132725352 | 136.154 | 36.57126935 | 0 |
| 4.44893E-07 | 0.169470657 | 8 | -1.641322354 | 0.354117793 | 14.17570281 | 6.560752194 | 0.4 |
| 4.48038E-07 | 0.004796206 | 91 | -2.212677756 | 0.159731975 | 231.579 | 51.24767393 | 0 |
| 4.52654E-07 | 0.186283576 | 46 | -1.552785915 | 0.393812293 | 12.62825651 | 6.178479099 | 0.2 |
| 4.56315E-07 | 0.181095092 | 36 | -1.590898111 | 0.399559542 | 13.17771084 | 6.001804183 | 0.4 |
| 4.56356E-07 | 0.199225753 | 72 | -1.476786266 | 0.423571001 | 11.66565962 | 5.78452111 | 0.7 |
| 4.57230E-07 | 0.151610343 | 43 | -1.732066618 | 0.384118326 | 17.00900901 | 7.852403415 | 0.1 |
| 4.57358E-07 | 0.187871408 | 27 | -1.534098368 | 0.422468046 | 12.70251256 | 6.327413428 | 0.5 |
| 4.58596E-07 | 0.116048413 | 48 | -1.961712174 | 0.280322732 | 25.57057057 | 10.3504124 | 0.1 |
| 4.59638E-07 | 0.053993528 | 73 | -2.300837062 | 0.154081865 | 59.535 | 20.63842823 | 0 |
| 4.65103E-07 | 0.116094108 | 28 | -1.962340759 | 0.291183615 | 25.40280561 | 10.54103469 | 0.2 |
| 4.66800E-07 | 0.161534222 | 85 | -1.695431883 | 0.380400702 | 16.03106212 | 7.387133836 | 0.2 |
| 4.72373E-07 | 0.105331015 | 61 | -2.04391141 | 0.258162119 | 29.021 | 11.77017825 | 0 |
| 4.73611E-07 | 0.013217151 | 99 | -2.338045263 | 0.130808871 | 167.561 | 42.67612366 | 0 |
| 4.74453E-07 | 0.075842681 | 41 | -2.201958167 | 0.183824769 | 41.642 | 15.26974135 | 0 |
| 4.76678E-07 | 1.81E-05 | 80 | -0.828796119 | 0.473864262 | 627.398 | 139.04378 | 0 |
| 4.78537E-07 | 0.179169857 | 19 | -1.606507287 | 0.406218549 | 14.59337349 | 6.715489952 | 0.4 |
| 4.81157E-07 | 0.160491168 | 58 | -1.72225111 | 0.350772212 | 16.67667668 | 7.844465427 | 0.1 |
| 4.85082E-07 | 0.137902666 | 4 | -1.857548224 | 0.324507283 | 20.85785786 | 9.237992033 | 0.1 |
| 4.85231E-07 | 0.183335963 | 55 | -1.609872234 | 0.373760088 | 14.39034205 | 6.840032868 | 0.6 |
| 4.86011E-07 | 0.131413846 | 12 | -1.881475385 | 0.307215953 | 22.098 | 9.195812034 | 0 |
| 4.86422E-07 | 0.022695217 | 3 | -2.373722617 | 0.125346109 | 123.777 | 35.01578753 | 0 |
| 4.87254E-07 | 0.01990165 | 26 | -2.36873971 | 0.123626619 | 134.026 | 38.07943497 | 0 |
| 4.89744E-07 | 0.098521639 | 81 | -2.077731826 | 0.223896387 | 31.116 | 12.53098438 | 0 |
| 4.92587E-07 | 0.198861697 | 51 | -1.479313208 | 0.433173479 | 12.18932528 | 5.880737319 | 0.7 |
| 4.93722E-07 | 0.04876874 | 3 | -2.325216077 | 0.153559391 | 69.615 | 24.17443232 | 0 |
| 4.99492E-07 | 0.158744752 | 85 | -1.723296886 | 0.371951938 | 17.81024096 | 7.74136806 | 0.4 |
| 5.07314E-07 | 0.12820882 | 74 | -1.93328967 | 0.278126762 | 23.854 | 9.985910194 | 0 |
| 5.08780E-07 | 0.172074038 | 94 | -1.68434778 | 0.381311733 | 16.54964895 | 7.634522652 | 0.3 |
| 5.09719E-07 | 0.002444161 | 36 | -2.099328143 | 0.158381459 | 305.533 | 62.92273183 | 0 |
| 5.10720E-07 | 0.187349958 | 84 | -1.584976378 | 0.400536473 | 14.19539078 | 6.911889862 | 0.2 |
| 5.12413E-07 | 0.014328371 | 75 | -2.353418201 | 0.129180266 | 167.381 | 43.14941148 | 0 |
| 5.22986E-07 | 0.027897215 | 39 | -2.376066156 | 0.128146985 | 113.429 | 32.94559715 | 0 |
| 5.26933E-07 | 0.018889169 | 53 | -2.374738531 | 0.129071437 | 147.882 | 39.6916049 | 0 |
| 5.30076E-07 | 0.086444759 | 47 | -2.170540461 | 0.210112043 | 41.69 | 15.81385399 | 0 |
| 5.31326E-07 | 0.092535834 | 63 | -2.119109716 | 0.239276734 | 37.306 | 14.97606913 | 0 |
| 5.41062E-07 | 0.142412853 | 61 | -1.854984828 | 0.323955412 | 22.439 | 9.4031217 | 0 |
| 5.41403E-07 | 0.098318418 | 61 | -2.117486568 | 0.220078354 | 36.471 | 13.97671304 | 0 |
| 5.45228E-07 | 0.071245859 | 56 | -2.241210892 | 0.187757874 | 50.269 | 18.60469763 | 0 |
| 5.46210E-07 | 0.089440989 | 94 | -2.161198011 | 0.211059968 | 40.233 | 14.83964036 | 0 |
| 5.47140E-07 | 0.18887474 | 83 | -1.591718244 | 0.40295329 | 15.03614458 | 6.949256046 | 0.4 |
| 5.50747E-07 | 0.092513615 | 87 | -2.132668265 | 0.227529601 | 38.957 | 15.1101114 | 0 |
| 5.55891E-07 | 0.041245357 | 79 | -2.358015556 | 0.140884001 | 87.064 | 28.91944395 | 0 |
| 5.62946E-07 | 0.127920307 | 1 | -1.935608969 | 0.301668399 | 26.14414414 | 10.81272738 | 0.1 |
| 5.63912E-07 | 0.061802322 | 41 | -2.286673673 | 0.176634746 | 61.096 | 21.66772425 | 0 |
| 5.67740E-07 | 0.073935846 | 64 | -2.241936708 | 0.17885454 | 50.633 | 18.87311595 | 0 |
| 5.73891E-07 | 0.036132675 | 51 | -2.381969483 | 0.127099659 | 100.507 | 31.59592147 | 0 |
| 5.76137E-07 | 0.093415537 | 69 | -2.135582306 | 0.233306332 | 40.19 | 15.81980297 | 0 |
| 5.77759E-07 | 0.136922281 | 1 | -1.891555179 | 0.317840469 | 25.21021021 | 10.53754885 | 0.1 |
| 5.83091E-07 | 0.118965916 | 71 | -2.009976284 | 0.275477983 | 30.95 | 12.67636345 | 0 |
| 5.86024E-07 | 0.042663324 | 28 | -2.365750061 | 0.144936209 | 91.262 | 29.73596448 | 0 |
| 5.87869E-07 | 0.068704733 | 30 | -2.268775213 | 0.174609937 | 56.323 | 20.37085277 | 0 |
| 5.88930E-07 | 0.151381812 | 81 | -1.835404395 | 0.321817992 | 22.28928929 | 9.451245817 | 0.1 |
| 5.89784E-07 | 0.121789582 | 5 | -1.985400331 | 0.287559606 | 29.799 | 11.78386433 | 0 |
| 5.92252E-07 | 0.012728503 | 89 | -2.365315467 | 0.126212424 | 195.285 | 48.73625958 | 0 |
| 5.97764E-07 | 0.061090506 | 26 | -2.300659524 | 0.164891714 | 64.284 | 22.72980396 | 0 |
| 6.07912E-07 | 0.064251841 | 3 | -2.292260897 | 0.170357909 | 63.575 | 23.29611736 | 0 |
| 6.10206E-07 | 0.188284389 | 45 | -1.627234139 | 0.419554958 | 16.91291291 | 7.974874335 | 0.1 |
| 6.11120E-07 | 0.133549017 | 28 | -1.934261205 | 0.301857247 | 27.205 | 11.16170052 | 0 |
| 6.22853E-07 | 0.161318926 | 63 | -1.786585267 | 0.359595657 | 21.63563564 | 9.413852166 | 0.1 |
| 6.25570E-07 | 0.105957438 | 85 | -2.08558759 | 0.258754248 | 37.698 | 15.06823971 | 0 |
| 6.27399E-07 | 0.021757202 | 56 | -2.39269476 | 0.124485746 | 150.373 | 41.96442403 | 0 |
| 6.29198E-07 | 0.068921262 | 90 | -2.268622756 | 0.182335149 | 59.777 | 21.2884586 | 0 |
| 6.31635E-07 | 0.10812276 | 39 | -2.074949967 | 0.271997634 | 37.18018018 | 14.75027827 | 0.1 |
| 6.32878E-07 | 0.126549014 | 23 | -1.967960893 | 0.29350933 | 29.778 | 12.50388192 | 0 |
| 6.33823E-07 | 0.178907936 | 73 | -1.671331584 | 0.389386425 | 18.62725451 | 8.126466455 | 0.2 |
| 6.34670E-07 | 0.154019599 | 35 | -1.821776121 | 0.367887564 | 23.32 | 10.25567057 | 0 |
| 6.41006E-07 | 0.003459179 | 72 | -2.195076763 | 0.160541996 | 332.256 | 69.47318178 | 0 |
| 6.46468E-07 | 0.185159712 | 30 | -1.67006804 | 0.386386813 | 18.021 | 8.141058643 | 0 |
| 6.48995E-07 | 0.073932404 | 98 | -2.257821508 | 0.188034544 | 57.549 | 21.1703515 | 0 |
| 6.49178E-07 | 0.005010425 | 31 | -2.26484028 | 0.133943974 | 298.541 | 65.20969731 | 0 |
| 6.49894E-07 | 0.11317488 | 43 | -2.057983013 | 0.253372833 | 35.365 | 14.2420016 | 0 |
| 6.51568E-07 | 0.063328497 | 14 | -2.306086825 | 0.170116923 | 68.583 | 24.40845927 | 0 |
| 6.54198E-07 | 0.174952414 | 5 | -1.735161693 | 0.36212374 | 19.96693387 | 9.099136865 | 0.2 |
| 6.56364E-07 | 0.06533232 | 43 | -2.309508619 | 0.163992778 | 66.057 | 22.59646767 | 0 |
| 6.56444E-07 | 0.12333492 | 71 | -2.010133434 | 0.259923441 | 32.09309309 | 13.0049452 | 0.1 |
| 6.57346E-07 | 0.056927165 | 57 | -2.33082492 | 0.157865604 | 74.947 | 26.47869544 | 0 |
| 6.63547E-07 | 0.027365637 | 8 | -2.395972024 | 0.129348737 | 137.284 | 40.18300658 | 0 |
| 6.73692E-07 | 0.067769351 | 19 | -2.282983501 | 0.176197349 | 64.356 | 22.75044718 | 0 |
| 6.74776E-07 | 0.196447317 | 43 | -1.613491562 | 0.40610367 | 17.39418255 | 8.310012726 | 0.3 |
| 6.78815E-07 | 0.17252288 | 2 | -1.739164739 | 0.386092426 | 20.58458458 | 8.968359133 | 0.1 |
| 6.90168E-07 | 0.065720653 | 72 | -2.303813215 | 0.185377957 | 68.618 | 24.48807346 | 0 |
| 6.93034E-07 | 0.172551217 | 87 | -1.726582668 | 0.414998201 | 21.269 | 9.367517782 | 0 |
| 7.00209E-07 | 0.112848824 | 13 | -2.072358438 | 0.262180829 | 38.486 | 15.41484419 | 0 |
| 7.09381E-07 | 0.160081838 | 6 | -1.815992771 | 0.367069638 | 24.058 | 10.77073092 | 0 |
| 7.15598E-07 | 0.165907051 | 83 | -1.796755044 | 0.35085572 | 23.47895792 | 10.46928248 | 0.2 |
| 7.18303E-07 | 0.172914761 | 12 | -1.752973161 | 0.37337729 | 22.842 | 10.02544491 | 0 |
| 7.23038E-07 | 0.04067889 | 20 | -2.399535512 | 0.134623914 | 109.438 | 35.41327797 | 0 |
| 7.23864E-07 | 0.03023938 | 10 | -2.413917369 | 0.122199938 | 139.245 | 40.93830627 | 0 |
| 7.27150E-07 | 0.068464767 | 65 | -2.305793295 | 0.173574978 | 68.654 | 24.55596219 | 0 |
| 7.28669E-07 | 0.170839024 | 21 | -1.782422594 | 0.346860099 | 22.904 | 10.1905167 | 0 |
| 7.32970E-07 | 0.029525725 | 27 | -2.411539847 | 0.123930862 | 140.514 | 41.50091722 | 0 |
| 7.35494E-07 | 0.109352402 | 57 | -2.129564421 | 0.262158095 | 42.374 | 16.45627336 | 0 |
| 7.37999E-07 | 0.064538844 | 81 | -2.312447822 | 0.166700053 | 74.775 | 26.43901277 | 0 |
| 7.38471E-07 | 0.08708861 | 88 | -2.217086612 | 0.20560124 | 53.985 | 20.49851126 | 0 |
| 7.40200E-07 | 0.007122198 | 86 | -2.31763108 | 0.131482564 | 287.851 | 66.02461699 | 0 |
| 7.43858E-07 | 0.11097448 | 35 | -2.093753998 | 0.262317469 | 40.736 | 16.4614484 | 0 |
| 7.45283E-07 | 0.170784282 | 98 | -1.769886522 | 0.3851698 | 23.35535536 | 9.907834234 | 0.1 |
| 7.45887E-07 | 0.133740643 | 95 | -1.963543268 | 0.307146824 | 32.626 | 13.65622996 | 0 |
| 7.46177E-07 | 0.11206069 | 49 | -2.089201067 | 0.270413636 | 40.621 | 16.06497564 | 0 |
| 7.48587E-07 | 0.186016318 | 11 | -1.686835983 | 0.397049695 | 20.669 | 9.039311265 | 0 |
| 7.54685E-07 | 0.050743669 | 12 | -2.368816153 | 0.143035592 | 94.978 | 31.31974525 | 0 |
| 7.56554E-07 | 0.083511943 | 5 | -2.225624981 | 0.214401708 | 58.178 | 21.59436576 | 0 |
| 7.57696E-07 | 0.118715895 | 99 | -2.052701247 | 0.273423057 | 37.426 | 14.9596228 | 0 |
| 7.66015E-07 | 0.103301166 | 9 | -2.142747128 | 0.236139051 | 46.152 | 17.97325607 | 0 |
| 7.67139E-07 | 0.122464078 | 44 | -2.039258757 | 0.295693826 | 38.223 | 15.98138529 | 0 |
| 7.70460E-07 | 0.124639136 | 50 | -2.020184672 | 0.289891967 | 36.467 | 14.2701278 | 0 |
| 7.73181E-07 | 0.109563451 | 56 | -2.124596041 | 0.265118401 | 43.723 | 16.6645125 | 0 |
| 7.73734E-07 | 0.124131001 | 5 | -2.039788628 | 0.275645329 | 37.339 | 14.8165316 | 0 |
| 7.78532E-07 | 0.157891765 | 88 | -1.866056127 | 0.31694272 | 27.212 | 11.84699993 | 0 |
| 7.81036E-07 | 0.113441656 | 88 | -2.104894273 | 0.242571695 | 42.251 | 17.24341423 | 0 |
| 7.83887E-07 | 0.071466447 | 8 | -2.300577909 | 0.174653556 | 70.637 | 25.09424141 | 0 |
| 7.87408E-07 | 0.186779623 | 60 | -1.687090532 | 0.417185199 | 21.37037037 | 9.599858203 | 0.1 |
| 7.88540E-07 | 0.119359156 | 87 | -2.073129193 | 0.268431024 | 40.545 | 16.40503277 | 0 |
| 7.91644E-07 | 0.165678778 | 3 | -1.809017429 | 0.369616887 | 26.101 | 11.59565687 | 0 |
| 7.91907E-07 | 0.009790842 | 64 | -2.362107843 | 0.122243822 | 263.733 | 62.03558757 | 0 |
| 7.95075E-07 | 0.056661637 | 65 | -2.359263741 | 0.147817899 | 88.329 | 30.44167299 | 0 |
| 8.02179E-07 | 0.118936536 | 93 | -2.065181003 | 0.277919237 | 40.489 | 16.40405352 | 0 |
| 8.03831E-07 | 0.037715098 | 27 | -2.408939246 | 0.126806834 | 124.749 | 38.86672898 | 0 |
| 8.03971E-07 | 0.126911162 | 96 | -2.038649188 | 0.293079635 | 37.673 | 14.9958977 | 0 |
| 8.11561E-07 | 0.057371531 | 47 | -2.359531602 | 0.157725104 | 90.97 | 30.8348486 | 0 |
| 8.23590E-07 | 0.047100156 | 17 | -2.392813941 | 0.136277983 | 110.282 | 34.90161312 | 0 |
| 8.24378E-07 | 0.126318938 | 38 | -2.04625646 | 0.276387324 | 38.651 | 15.02027988 | 0 |
| 8.26177E-07 | 0.119302398 | 1 | -2.073597824 | 0.274128725 | 41.908 | 16.18287433 | 0 |
| 8.31952E-07 | 0.146472046 | 42 | -1.952159634 | 0.307756778 | 32.588 | 14.11082462 | 0 |
| 8.33841E-07 | 0.096896669 | 78 | -2.202043903 | 0.214839821 | 54.368 | 20.76935102 | 0 |
| 8.38476E-07 | 0.002551945 | 68 | -2.226928671 | 0.141982707 | 390.475 | 83.64899599 | 0 |
| 8.42650E-07 | 0.127731742 | 62 | -2.035720753 | 0.29161473 | 38.438 | 15.94115041 | 0 |
| 8.42714E-07 | 0.001324957 | 32 | -2.079184 | 0.160879099 | 458.899 | 98.29746279 | 0 |
| 8.43452E-07 | 0.09768117 | 61 | -2.173849678 | 0.233342268 | 53.819 | 20.21323363 | 0 |
| 8.43739E-07 | 0.067761497 | 6 | -2.32000126 | 0.176019352 | 78.947 | 28.20400072 | 0 |
| 8.45572E-07 | 0.096051999 | 25 | -2.178944854 | 0.231792045 | 54.742 | 21.13433453 | 0 |
| 8.45649E-07 | 0.183461131 | 6 | -1.729375924 | 0.383622783 | 24.14714715 | 10.73796896 | 0.1 |
| 8.46633E-07 | 0.087285589 | 4 | -2.223955615 | 0.221679003 | 59.829 | 22.29041547 | 0 |
| 8.47860E-07 | 0.10837639 | 53 | -2.122915155 | 0.251857677 | 48.35 | 18.69521359 | 0 |
| 8.50572E-07 | 0.151922227 | 37 | -1.892116848 | 0.340573934 | 31.512 | 13.80660996 | 0 |
| 8.58158E-07 | 0.041572711 | 27 | -2.412564388 | 0.13991195 | 122.952 | 37.92827771 | 0 |
| 8.60427E-07 | 0.076306496 | 11 | -2.29049708 | 0.179318653 | 70.689 | 24.50376938 | 0 |
| 8.63109E-07 | 0.018727661 | 82 | -2.41929724 | 0.11599439 | 209.869 | 57.66908672 | 0 |
| 8.67748E-07 | 0.013888134 | 43 | -2.400844977 | 0.118750513 | 242.481 | 59.70387082 | 0 |
| 8.72738E-07 | 0.151899147 | 57 | -1.908351221 | 0.333381894 | 32.321 | 13.70298255 | 0 |
| 8.79088E-07 | 0.188572366 | 20 | -1.731098666 | 0.392420045 | 23.61661662 | 10.46514226 | 0.1 |
| 8.90965E-07 | 0.076500045 | 30 | -2.279598181 | 0.19632722 | 72.923 | 26.49900504 | 0 |
| 8.93050E-07 | 0.189576839 | 94 | -1.721203809 | 0.426115355 | 23.83 | 10.84853865 | 0 |
| 8.94638E-07 | 0.088764927 | 77 | -2.229402964 | 0.213832092 | 61.753 | 23.17703963 | 0 |
| 8.95009E-07 | 0.168161572 | 30 | -1.821728785 | 0.354682282 | 28.7987988 | 12.21896742 | 0.1 |
| 8.95196E-07 | 0.071280684 | 69 | -2.31169661 | 0.168400319 | 79.715 | 27.37299867 | 0 |
| 1.02549E-08 | 0.166621462 | 95 | -0.927770559 | 0.303715992 | 2.095238095 | 0.297101757 | 95.8 |
| 1.06950E-08 | 0.040927772 | 48 | -1.258643589 | 0.237590368 | 3.196638655 | 1.430778365 | 40.5 |
| 1.08510E-08 | 0.044886013 | 80 | -1.257029637 | 0.211540237 | 3.107407407 | 1.239254646 | 46 |
| 1.10037E-08 | 0.027449551 | 44 | -1.349544288 | 0.264949911 | 4.158653846 | 2.006325044 | 16.8 |
| 1.10591E-08 | 0.18920789 | 62 | -0.989228572 | 0.329183214 | 2.136363636 | 0.408679615 | 95.6 |
| 1.13372E-08 | 0.071016741 | 54 | -1.176759065 | 0.19201822 | 2.513368984 | 0.791091748 | 62.6 |
| 1.13524E-08 | 0.100426938 | 67 | -1.116350232 | 0.214595091 | 2.381188119 | 0.682578207 | 79.8 |
| 1.18474E-08 | 0.186101365 | 97 | -0.938338761 | 0.276544438 | 2.055555556 | 0.231212282 | 94.6 |
| 1.19672E-08 | 0.025621418 | 80 | -1.374937961 | 0.260214865 | 4.498281787 | 2.085953301 | 12.7 |
| 1.31692E-08 | 0.019304788 | 75 | -1.47088918 | 0.276639827 | 6.058762887 | 2.82982248 | 3 |
| 1.37567E-08 | 0.191421556 | 45 | -0.891193972 | 0.327302406 | 2.229508197 | 0.588598409 | 93.9 |
| 1.38273E-08 | 0.002268162 | 93 | -1.232773757 | 0.461863392 | 19.673 | 6.084002473 | 0 |
| 1.38322E-08 | 0.004473975 | 55 | -1.398755614 | 0.367019519 | 15.665 | 5.113210334 | 0 |
| 1.39104E-08 | 0.097980034 | 45 | -1.135839726 | 0.220761062 | 2.526717557 | 0.725329829 | 73.8 |
| 1.42461E-08 | 0.157215948 | 94 | -0.955310389 | 0.294724987 | 2.2 | 0.447213595 | 89.5 |
| 1.49193E-08 | 0.032895211 | 3 | -1.393375055 | 0.259908002 | 4.393764434 | 2.127368134 | 13.4 |
| 1.50476E-08 | 0.174141262 | 35 | -1.00275352 | 0.235382276 | 2.295238095 | 0.517460692 | 89.5 |
| 1.51772E-08 | 0.178423798 | 65 | -0.950406587 | 0.317505773 | 2.2 | 0.451946146 | 90.5 |
| 1.52825E-08 | 0.160833498 | 96 | -0.988483119 | 0.33707976 | 2.170212766 | 0.377834744 | 90.6 |
| 1.58587E-08 | 0.028057542 | 54 | -1.460617053 | 0.265502247 | 5.25958379 | 2.494177132 | 8.7 |
| 1.63525E-08 | 0.104537563 | 51 | -1.136497959 | 0.230661079 | 2.528239203 | 0.814473177 | 69.9 |
| 1.70225E-08 | 0.134994325 | 99 | -1.063572288 | 0.257651468 | 2.380952381 | 0.745808959 | 81.1 |
| 1.72272E-08 | 0.162494036 | 91 | -1.005188644 | 0.299550167 | 2.276785714 | 0.54043395 | 88.8 |
| 1.77815E-08 | 0.06838154 | 25 | -1.235681601 | 0.225890137 | 2.978533095 | 1.189667582 | 44.1 |
| 1.82548E-08 | 0.001912839 | 99 | -1.309060885 | 0.42393027 | 24.852 | 7.278122496 | 0 |
| 1.82589E-08 | 0.194236652 | 81 | -0.965604974 | 0.336085499 | 2.220930233 | 0.470323113 | 91.4 |
| 1.86388E-08 | 0.168700576 | 34 | -1.019028473 | 0.267958198 | 2.345588235 | 0.6131724 | 86.4 |
| 1.87695E-08 | 0.04253625 | 20 | -1.397551404 | 0.247532211 | 4.240143369 | 1.922770623 | 16.3 |
| 1.93798E-08 | 0.029317604 | 67 | -1.518148596 | 0.267990532 | 6.057894737 | 2.760517924 | 5 |
| 2.00498E-08 | 0.147561911 | 2 | -1.041229389 | 0.250834505 | 2.398989899 | 0.717722703 | 80.2 |
| 2.01275E-08 | 0.119226668 | 90 | -1.107970939 | 0.260777195 | 2.53125 | 0.799073719 | 68 |
| 2.06463E-08 | 0.033862157 | 66 | -1.498791879 | 0.281356725 | 5.629669157 | 2.613756997 | 6.3 |
| 2.07996E-08 | 0.08871058 | 37 | -1.20025501 | 0.230868452 | 2.815286624 | 1.036493615 | 52.9 |
| 2.08299E-08 | 0.044935262 | 79 | -1.40027667 | 0.258072736 | 4.409038239 | 2.135928837 | 13.7 |
| 2.10308E-08 | 0.188900915 | 92 | -0.930119822 | 0.313707961 | 2.295454545 | 0.534870126 | 86.8 |
| 2.13643E-08 | 0.186701684 | 98 | -1.022212528 | 0.290788563 | 2.348484848 | 0.641858567 | 86.8 |
| 2.17089E-08 | 0.126026928 | 68 | -1.097393858 | 0.273962162 | 2.605015674 | 0.961903209 | 68.1 |
| 2.17641E-08 | 0.019118405 | 96 | -1.661553305 | 0.260471933 | 9.518108652 | 4.004299558 | 0.6 |
| 2.19014E-08 | 0.088611417 | 59 | -1.19748045 | 0.239604912 | 2.880324544 | 1.106098347 | 50.7 |
| 2.27139E-08 | 0.04778282 | 6 | -1.411357093 | 0.251557644 | 4.427710843 | 2.095190267 | 17 |
| 2.28218E-08 | 0.193667433 | 13 | -0.96211986 | 0.30454337 | 2.201680672 | 0.461754795 | 88.1 |
| 2.33544E-08 | 0.190849318 | 66 | -0.977355022 | 0.313655215 | 2.322368421 | 0.636650732 | 84.8 |
| 2.36558E-08 | 0.001948288 | 88 | -1.374743654 | 0.411292592 | 30.966 | 8.485803018 | 0 |
| 2.42326E-08 | 0.108126355 | 4 | -1.159389608 | 0.245092304 | 2.777272727 | 1.088532823 | 56 |
| 2.42992E-08 | 0.142113581 | 61 | -1.065484424 | 0.233305221 | 2.443661972 | 0.728274478 | 71.6 |
| 2.43615E-08 | 0.158688121 | 81 | -1.068279613 | 0.279633649 | 2.517073171 | 0.770759251 | 79.5 |
| 2.43992E-08 | 0.00658896 | 63 | -1.694715431 | 0.28573051 | 20.76 | 6.63041193 | 0 |
| 2.49798E-08 | 0.010398879 | 45 | -1.748196946 | 0.265568396 | 17.148 | 6.033780117 | 0 |
| 2.51393E-08 | 0.186037772 | 52 | -0.984606054 | 0.315367101 | 2.357575758 | 0.68031122 | 83.5 |
| 2.52103E-08 | 0.126946612 | 66 | -1.122561726 | 0.261626974 | 2.653521127 | 0.930316602 | 64.5 |
| 2.59100E-08 | 0.116103656 | 74 | -1.112509918 | 0.269557827 | 2.72361809 | 1.025614014 | 60.2 |
| 2.61081E-08 | 0.162699277 | 46 | -1.048846153 | 0.247566932 | 2.408071749 | 0.728646378 | 77.7 |
| 2.63367E-08 | 0.077065189 | 13 | -1.294867779 | 0.253340497 | 3.54294032 | 1.536002944 | 31.3 |
| 2.63809E-08 | 0.06712947 | 27 | -1.327597687 | 0.257203152 | 3.785620915 | 1.771960871 | 23.5 |
| 2.69864E-08 | 0.196584834 | 30 | -0.947872541 | 0.320109436 | 2.347058824 | 0.664053936 | 83 |
| 2.71456E-08 | 0.193349643 | 16 | -0.946533214 | 0.339147232 | 2.388888889 | 0.654924525 | 82 |
| 2.72271E-08 | 0.179197088 | 60 | -0.99816553 | 0.30528566 | 2.391509434 | 0.690163512 | 78.8 |
| 2.72913E-08 | 0.051432378 | 32 | -1.453762469 | 0.260203136 | 4.895761741 | 2.237994043 | 12.7 |
| 2.74122E-08 | 0.126417189 | 93 | -1.119262301 | 0.273767692 | 2.722527473 | 1.037818323 | 63.6 |
| 2.74777E-08 | 0.187174034 | 43 | -0.939425824 | 0.344707126 | 2.434782609 | 0.772538896 | 81.6 |
| 2.77261E-08 | 0.135177669 | 48 | -1.096222267 | 0.274659883 | 2.626740947 | 0.933521127 | 64.1 |
| 2.77473E-08 | 0.011572525 | 50 | -1.787263772 | 0.245301687 | 16.83583584 | 5.907187724 | 0.1 |
| 2.78436E-08 | 0.158181546 | 29 | -1.075676761 | 0.28069892 | 2.625 | 0.914344911 | 73.6 |
| 2.79716E-08 | 0.121370404 | 23 | -1.140570717 | 0.22730336 | 2.767494357 | 1.066830124 | 55.7 |
| 2.80729E-08 | 0.124909106 | 73 | -1.119453462 | 0.278057272 | 2.762626263 | 1.071715611 | 60.4 |
| 2.82604E-08 | 0.181244541 | 30 | -0.979477082 | 0.321192902 | 2.418719212 | 0.729196177 | 79.7 |
| 2.86772E-08 | 0.018544266 | 96 | -1.767353907 | 0.241837272 | 12.40540541 | 4.853444412 | 0.1 |
| 2.93195E-08 | 0.130182068 | 74 | -1.09413657 | 0.290354767 | 2.743529412 | 1.005931355 | 57.5 |
| 3.01743E-08 | 0.18749573 | 90 | -0.984159067 | 0.285103658 | 2.372641509 | 0.658741155 | 78.8 |
| 3.03145E-08 | 0.038624241 | 15 | -1.591359269 | 0.253253933 | 6.70880829 | 3.20256237 | 3.5 |
| 3.03352E-08 | 0.072671652 | 78 | -1.313701349 | 0.262304542 | 3.765306122 | 1.79996322 | 21.6 |
| 3.04411E-08 | 0.068350278 | 19 | -1.356203388 | 0.2564287 | 4.048689139 | 1.92779055 | 19.9 |
| 3.10588E-08 | 0.024551968 | 14 | -1.740643367 | 0.245955077 | 10.40080564 | 4.45321551 | 0.7 |
| 3.10593E-08 | 0.109796989 | 53 | -1.173573286 | 0.272641215 | 3.001988072 | 1.264752005 | 49.7 |
| 3.15457E-08 | 0.006028588 | 66 | -1.743420009 | 0.278227625 | 27.554 | 7.935619576 | 0 |
| 3.16380E-08 | 0.03873084 | 24 | -1.599565843 | 0.25658156 | 6.845435685 | 3.172108771 | 3.6 |
| 3.19440E-08 | 0.126001308 | 32 | -1.150032042 | 0.257219056 | 2.902953586 | 1.138910002 | 52.6 |
| 3.28659E-08 | 0.188181744 | 55 | -1.012516885 | 0.290534947 | 2.445544554 | 0.785126 | 79.8 |
| 3.28880E-08 | 0.070431639 | 45 | -1.380510201 | 0.255350269 | 4.323636364 | 2.009863146 | 17.5 |
| 3.29551E-08 | 0.128145353 | 22 | -1.154887402 | 0.281660236 | 2.969565217 | 1.229249477 | 54 |
| 3.32757E-08 | 0.15603905 | 62 | -1.056789878 | 0.289017561 | 2.611594203 | 0.964415752 | 65.5 |
| 3.34054E-08 | 0.131182663 | 33 | -1.142894011 | 0.298182441 | 2.887417219 | 1.178023579 | 54.7 |
| 3.40177E-08 | 0.075036148 | 81 | -1.338646579 | 0.271325031 | 4.005025126 | 1.949029721 | 20.4 |
| 3.46501E-08 | 0.037579753 | 47 | -1.641431633 | 0.262216759 | 7.686734694 | 3.578797447 | 2 |
| 3.54171E-08 | 0.13890189 | 18 | -1.106733977 | 0.270906155 | 2.764150943 | 1.065707785 | 57.6 |
| 3.58636E-08 | 0.03356998 | 77 | -1.707681451 | 0.256625339 | 9.06612411 | 3.989127527 | 1.7 |
| 3.60594E-08 | 0.077295303 | 63 | -1.360822456 | 0.254736297 | 4.15776699 | 1.963051872 | 17.6 |
| 3.64318E-08 | 0.135068725 | 40 | -1.134055041 | 0.283041339 | 2.919354839 | 1.183023172 | 50.4 |
| 3.67460E-08 | 0.088118733 | 24 | -1.306619464 | 0.277618495 | 3.779548473 | 1.792198314 | 24.7 |
| 3.74177E-08 | 0.085571195 | 24 | -1.314211951 | 0.288079899 | 3.898832685 | 1.792520155 | 22.9 |
| 3.79671E-08 | 0.097927423 | 31 | -1.277687544 | 0.280236214 | 3.627737226 | 1.626746921 | 31.5 |
| 3.83399E-08 | 0.18967709 | 84 | -0.948242067 | 0.347297603 | 2.518394649 | 0.8286984 | 70.1 |
| 3.87822E-08 | 0.160174614 | 45 | -1.07006409 | 0.3153692 | 2.712328767 | 0.97326137 | 63.5 |
| 3.88599E-08 | 0.074314286 | 71 | -1.392087492 | 0.279256515 | 4.544273908 | 2.124964116 | 15.3 |
| 3.97460E-08 | 0.121794102 | 59 | -1.176261446 | 0.263746461 | 3.086805556 | 1.291887545 | 42.4 |
| 3.98361E-08 | 0.110882519 | 14 | -1.223632301 | 0.272236659 | 3.368499257 | 1.520990134 | 32.7 |
| 4.00437E-08 | 0.007419299 | 65 | -1.846489668 | 0.252321243 | 31.474 | 8.789230602 | 0 |
| 4.00787E-08 | 0.179253021 | 81 | -1.034597282 | 0.316023444 | 2.618902439 | 0.897435703 | 67.2 |
| 4.06806E-08 | 0.004447736 | 33 | -1.728623338 | 0.282180432 | 40.536 | 10.68198525 | 0 |
| 4.10283E-08 | 0.179458796 | 67 | -1.035383639 | 0.326309415 | 2.576923077 | 0.841258066 | 66.2 |
| 4.17174E-08 | 0.064628208 | 43 | -1.488539862 | 0.278977912 | 5.521546961 | 2.765109304 | 9.5 |
| 4.18112E-08 | 0.014176559 | 58 | -1.911656544 | 0.215369676 | 22.101 | 7.368520482 | 0 |
| 4.21990E-08 | 0.069601105 | 87 | -1.448272612 | 0.273257538 | 5.067114094 | 2.59510547 | 10.6 |
| 4.22649E-08 | 0.059863422 | 3 | -1.52545127 | 0.273099346 | 5.825531915 | 2.751587253 | 6 |
| 4.22991E-08 | 0.068692159 | 83 | -1.464700005 | 0.271343289 | 5.1171875 | 2.455570639 | 10.4 |
| 4.30696E-08 | 0.093657618 | 49 | -1.319643736 | 0.268975063 | 4.039042821 | 2.047909669 | 20.6 |
| 4.30767E-08 | 0.120789975 | 19 | -1.204555362 | 0.26835306 | 3.321821036 | 1.396889338 | 36.3 |
| 4.40754E-08 | 0.150367728 | 4 | -1.116256619 | 0.290956678 | 3.004106776 | 1.247212354 | 51.3 |
| 4.41865E-08 | 0.174184304 | 47 | -1.065990165 | 0.309784824 | 2.800970874 | 1.091528551 | 58.8 |
| 4.42550E-08 | 0.048952082 | 28 | -1.632507696 | 0.271695922 | 7.392820513 | 3.559489983 | 2.5 |
| 4.45005E-08 | 0.00605235 | 67 | -1.838727553 | 0.262381873 | 37.079 | 10.21661089 | 0 |
| 4.51102E-08 | 0.116337226 | 30 | -1.225835771 | 0.278870589 | 3.515151515 | 1.584661151 | 34 |
| 4.51624E-08 | 0.008590777 | 67 | -1.902690269 | 0.239763927 | 31.809 | 9.51015047 | 0 |
| 4.52507E-08 | 0.072360345 | 27 | -1.473201546 | 0.27838139 | 5.336225597 | 2.591175634 | 7.8 |
| 4.52644E-08 | 0.049148661 | 7 | -1.640454573 | 0.275062185 | 7.625514403 | 3.512474055 | 2.8 |
| 4.58375E-08 | 0.082761083 | 11 | -1.390266488 | 0.263988306 | 4.568469505 | 2.192145611 | 13.1 |
| 4.59814E-08 | 0.015692463 | 76 | -1.939526346 | 0.21191459 | 22.119 | 7.431693553 | 0 |
| 4.65201E-08 | 0.087850088 | 88 | -1.375365351 | 0.274626443 | 4.430094787 | 2.148926072 | 15.6 |
| 4.69712E-08 | 0.169363729 | 77 | -1.056946782 | 0.316878342 | 2.922727273 | 1.247660889 | 56 |
| 4.74198E-08 | 0.15336794 | 44 | -1.101334501 | 0.312214703 | 3.005882353 | 1.361108956 | 49 |
| 4.77424E-08 | 0.141501597 | 52 | -1.139694211 | 0.331548142 | 3.19343696 | 1.455412913 | 42.1 |
| 4.77553E-08 | 0.1518446 | 82 | -1.121040941 | 0.306511403 | 3.082568807 | 1.282865304 | 45.5 |
| 4.79167E-08 | 0.179497246 | 48 | -1.054439002 | 0.304803988 | 2.708860759 | 0.994297175 | 60.5 |
| 4.80844E-08 | 0.145199405 | 61 | -1.115385886 | 0.309429088 | 3.070143885 | 1.322204267 | 44.4 |
| 4.93696E-08 | 0.056948265 | 49 | -1.605599545 | 0.273239165 | 7.047131148 | 3.42147546 | 2.4 |
| 4.95222E-08 | 0.125806707 | 84 | -1.213863937 | 0.259113039 | 3.459541985 | 1.480080719 | 34.5 |
| 4.95710E-08 | 0.121119899 | 73 | -1.232531223 | 0.291127904 | 3.631954351 | 1.657469768 | 29.9 |
| 4.97218E-08 | 0.106983586 | 29 | -1.276708714 | 0.311563474 | 3.909214092 | 1.897980554 | 26.2 |
| 5.00838E-08 | 0.01152962 | 31 | -1.965089939 | 0.211308235 | 28.862 | 8.914730979 | 0 |
| 5.11168E-08 | 0.168721148 | 63 | -1.068644978 | 0.331325362 | 2.915019763 | 1.152995109 | 49.4 |
| 5.14290E-08 | 0.163176623 | 86 | -1.104035862 | 0.296823345 | 3.005928854 | 1.233992324 | 49.4 |
| 5.15830E-08 | 0.087953779 | 20 | -1.376949878 | 0.286049984 | 4.653758542 | 2.228339958 | 12.2 |
| 5.17548E-08 | 0.161152842 | 41 | -1.090584051 | 0.297859921 | 3.024048096 | 1.187037945 | 50.1 |
| 5.19361E-08 | 0.062704661 | 41 | -1.595235379 | 0.282411658 | 6.860294118 | 3.356243944 | 4.8 |
| 5.28199E-08 | 0.042245061 | 71 | -1.779328133 | 0.247690781 | 10.45975855 | 4.487746587 | 0.6 |
| 5.28255E-08 | 0.063306001 | 46 | -1.567188051 | 0.281668395 | 6.61105318 | 3.207168842 | 4.1 |
| 5.31820E-08 | 0.008935906 | 60 | -1.948120994 | 0.231650327 | 36.118 | 10.16047791 | 0 |
| 5.33833E-08 | 0.196059525 | 42 | -0.9888226 | 0.339468613 | 2.664160401 | 0.949738782 | 60.1 |
| 5.36643E-08 | 0.106108984 | 82 | -1.321821568 | 0.283392556 | 4.155927835 | 1.948409685 | 22.4 |
| 5.37567E-08 | 0.115271647 | 96 | -1.262042197 | 0.291454253 | 3.822768435 | 1.731571838 | 22.7 |
| 5.37605E-08 | 0.136818281 | 26 | -1.174428789 | 0.308828459 | 3.341732283 | 1.451010794 | 36.5 |
| 5.42478E-08 | 0.007267228 | 87 | -1.902644506 | 0.239456324 | 40.917 | 10.80772492 | 0 |
| 5.45313E-08 | 0.122548245 | 11 | -1.235431792 | 0.279289648 | 3.67798913 | 1.622412508 | 26.4 |
| 5.47603E-08 | 0.109345728 | 20 | -1.299795208 | 0.296287224 | 4.131611316 | 1.954821496 | 18.7 |
| 5.56244E-08 | 0.10416775 | 53 | -1.334503008 | 0.291461337 | 4.369512195 | 2.121187323 | 18 |
| 5.56793E-08 | 0.12543903 | 51 | -1.226467123 | 0.303297667 | 3.671724138 | 1.825944978 | 27.5 |
| 5.57216E-08 | 0.180434003 | 80 | -1.082790968 | 0.314082881 | 3.078947368 | 1.257771825 | 50.6 |
| 5.58493E-08 | 0.193865959 | 78 | -1.0217415 | 0.347692376 | 2.826506024 | 1.076212308 | 58.5 |
| 5.60914E-08 | 0.058138095 | 29 | -1.667646791 | 0.264036245 | 8.047082907 | 3.731489037 | 2.3 |
| 5.61151E-08 | 0.112464925 | 61 | -1.305162748 | 0.286586333 | 4.119791667 | 1.918475046 | 23.2 |
| 5.64214E-08 | 0.195214354 | 54 | -1.000697513 | 0.322795056 | 2.84057971 | 1.106308414 | 58.6 |
| 5.64275E-08 | 0.187407629 | 14 | -1.026662064 | 0.346304436 | 2.915367483 | 1.148352823 | 55.1 |
| 5.66711E-08 | 0.18264967 | 63 | -1.037419448 | 0.321467929 | 2.884090909 | 1.18368622 | 56 |
| 5.69467E-08 | 0.095471112 | 95 | -1.386375075 | 0.297169207 | 4.825934579 | 2.399577022 | 14.4 |
| 5.77445E-08 | 0.035818002 | 73 | -1.874570294 | 0.223807816 | 13.02802803 | 5.214940693 | 0.1 |
| 5.80649E-08 | 0.146920977 | 22 | -1.163898799 | 0.314599189 | 3.381789137 | 1.532450271 | 37.4 |
| 5.82768E-08 | 0.13321176 | 78 | -1.212145201 | 0.291306829 | 3.552407932 | 1.649655415 | 29.4 |
| 5.83021E-08 | 0.051742495 | 9 | -1.716831162 | 0.264563631 | 9.013184584 | 4.178119048 | 1.4 |
| 5.88793E-08 | 0.025214899 | 55 | -1.97655638 | 0.209226376 | 19.125 | 6.973378163 | 0 |
| 5.97397E-08 | 0.151678629 | 64 | -1.141115748 | 0.323160391 | 3.373417722 | 1.486267524 | 36.8 |
| 5.98344E-08 | 0.041260736 | 11 | -1.832199046 | 0.245698481 | 11.90562249 | 4.958838464 | 0.4 |
| 6.04012E-08 | 0.151567184 | 32 | -1.157430252 | 0.290811915 | 3.322033898 | 1.421665969 | 35.1 |
| 6.04017E-08 | 0.046380679 | 38 | -1.791959762 | 0.258017848 | 10.77194753 | 4.819411522 | 0.9 |
| 6.04703E-08 | 0.171901895 | 29 | -1.113354796 | 0.300565883 | 3.130208333 | 1.256104658 | 42.4 |
| 6.08165E-08 | 0.078767198 | 58 | -1.514779552 | 0.29173394 | 6.097430407 | 3.021915988 | 6.6 |
| 6.08671E-08 | 0.080094909 | 40 | -1.51341542 | 0.290154984 | 6.069222577 | 2.962541649 | 6.1 |
| 6.09768E-08 | 0.04038015 | 95 | -1.858603574 | 0.235942864 | 12.66566567 | 5.250282208 | 0.1 |
| 6.16821E-08 | 0.093311023 | 70 | -1.413053499 | 0.305851606 | 5.192784667 | 2.539092509 | 11.3 |
| 6.18358E-08 | 0.003105391 | 23 | -1.757031883 | 0.284076314 | 61.184 | 15.78928261 | 0 |
| 6.19557E-08 | 0.000680834 | 10 | -1.472204287 | 0.370760764 | 78.733 | 18.19947614 | 0 |
| 6.25196E-08 | 0.043548086 | 61 | -1.847056048 | 0.240738027 | 11.91247485 | 5.026354042 | 0.6 |
| 6.26388E-08 | 0.006197232 | 3 | -1.910359979 | 0.241603025 | 49.656 | 12.78542651 | 0 |
| 6.28144E-08 | 0.089991688 | 42 | -1.430386712 | 0.287536753 | 5.266015201 | 2.545205697 | 7.9 |
| 6.29269E-08 | 0.062816499 | 28 | -1.655222565 | 0.266685946 | 8.060762101 | 3.726115005 | 2.9 |
| 6.29490E-08 | 0.115705576 | 38 | -1.299440858 | 0.300059441 | 4.261239368 | 2.044169662 | 17.7 |
| 6.30087E-08 | 0.02833647 | 39 | -1.97211339 | 0.213786015 | 18.08508509 | 6.827077394 | 0.1 |
| 6.32787E-08 | 0.11565336 | 8 | -1.298228507 | 0.300252073 | 4.216381418 | 2.03539683 | 18.2 |
| 6.34922E-08 | 0.177816587 | 63 | -1.065888142 | 0.327278154 | 3.118299445 | 1.308397971 | 45.9 |
| 6.37770E-08 | 0.091894077 | 37 | -1.425144197 | 0.297130513 | 5.27795874 | 2.696382956 | 7.9 |
| 6.39081E-08 | 0.136685745 | 87 | -1.213213947 | 0.30062682 | 3.742587601 | 1.715152064 | 25.8 |
| 6.44031E-08 | 0.062268 | 87 | -1.675588309 | 0.262221322 | 8.286004057 | 3.841708965 | 1.4 |
| 6.51204E-08 | 0.182882459 | 9 | -1.017862616 | 0.369157437 | 3.005545287 | 1.197979289 | 45.9 |
| 6.52500E-08 | 0.055291347 | 46 | -1.729548834 | 0.276368097 | 9.520812183 | 4.549839557 | 1.5 |
| 6.54777E-08 | 0.018066283 | 90 | -2.031396988 | 0.192812536 | 27.293 | 8.980857799 | 0 |
| 6.59931E-08 | 0.107429777 | 38 | -1.368804037 | 0.30531698 | 4.791860465 | 2.426028585 | 14 |
| 6.62527E-08 | 0.103102162 | 94 | -1.380413344 | 0.306068572 | 4.863945578 | 2.449414086 | 11.8 |
| 6.63110E-08 | 0.168995134 | 48 | -1.119446729 | 0.314576364 | 3.191558442 | 1.337025034 | 38.4 |
| 6.65924E-08 | 0.156218456 | 91 | -1.14480705 | 0.32362419 | 3.484210526 | 1.569707694 | 33.5 |
| 6.70064E-08 | 0.107908052 | 22 | -1.351492211 | 0.308453316 | 4.710801394 | 2.351889282 | 13.9 |
| 6.78114E-08 | 0.139843505 | 17 | -1.212771296 | 0.323831975 | 3.74796748 | 1.682023218 | 26.2 |
| 6.78407E-08 | 0.172280952 | 39 | -1.095421821 | 0.366439467 | 3.274873524 | 1.420289521 | 40.7 |
| 6.84194E-08 | 0.006145635 | 27 | -1.946685872 | 0.236363442 | 54.12 | 13.78069329 | 0 |
| 6.84434E-08 | 0.12504925 | 92 | -1.292133108 | 0.312171698 | 4.207286432 | 2.007782888 | 20.4 |
| 6.87551E-08 | 0.156191324 | 37 | -1.119010364 | 0.334869372 | 3.391629297 | 1.525920319 | 33.1 |
| 6.88514E-08 | 0.046376509 | 52 | -1.836189957 | 0.241212578 | 12.01603206 | 5.183662272 | 0.2 |
| 6.89897E-08 | 0.109093 | 83 | -1.358025461 | 0.312075186 | 4.790993072 | 2.357908779 | 13.4 |
| 6.90035E-08 | 0.038231201 | 30 | -1.906839032 | 0.238527403 | 14.6002004 | 5.859392237 | 0.2 |
| 7.10289E-08 | 0.011504202 | 9 | -2.037323738 | 0.200623867 | 38.597 | 11.53002743 | 0 |
| 7.16371E-08 | 0.119795949 | 33 | -1.316158395 | 0.313163254 | 4.464670659 | 2.129981194 | 16.5 |
| 7.19062E-08 | 0.108944477 | 12 | -1.380574391 | 0.288661169 | 4.992081448 | 2.469965582 | 11.6 |
| 7.21496E-08 | 0.098247163 | 75 | -1.436293783 | 0.308634183 | 5.46961326 | 2.786546052 | 9.5 |
| 7.25245E-08 | 0.103848526 | 67 | -1.426490338 | 0.302543919 | 5.431263858 | 2.54574973 | 9.8 |
| 7.27304E-08 | 0.117376649 | 22 | -1.349979489 | 0.289068135 | 4.754022989 | 2.19250762 | 13 |
| 7.32176E-08 | 0.146257167 | 77 | -1.200394421 | 0.32146923 | 3.733695652 | 1.815205746 | 26.4 |
| 7.39357E-08 | 0.067156525 | 6 | -1.681634223 | 0.279895496 | 8.593462717 | 4.078749127 | 2.1 |
| 7.41321E-08 | 0.053917111 | 80 | -1.797485041 | 0.250298879 | 10.92978937 | 4.867633994 | 0.3 |
| 7.47834E-08 | 0.01056615 | 62 | -2.041916749 | 0.197098565 | 42.509 | 12.23558861 | 0 |
| 7.53146E-08 | 0.035572741 | 85 | -1.977216193 | 0.209541238 | 17.444 | 6.943707877 | 0 |
| 7.57802E-08 | 0.10716773 | 18 | -1.406796573 | 0.31238153 | 5.28443449 | 2.702712812 | 10.7 |
| 7.64065E-08 | 0.138503474 | 65 | -1.240604605 | 0.331338091 | 4.090676884 | 2.014195765 | 21.7 |
| 7.64381E-08 | 0.196662457 | 81 | -1.038466995 | 0.350915754 | 3.114840989 | 1.288159375 | 43.4 |
| 7.64884E-08 | 0.104148309 | 41 | -1.437612716 | 0.291200144 | 5.499454744 | 2.621970018 | 8.3 |
| 7.66837E-08 | 0.035334821 | 34 | -1.982120303 | 0.207827013 | 17.65465465 | 6.702514697 | 0.1 |
| 7.70067E-08 | 0.104346935 | 4 | -1.434300125 | 0.305645228 | 5.468271335 | 2.664845554 | 8.6 |
| 7.70446E-08 | 0.119738701 | 48 | -1.347977488 | 0.287615347 | 4.806146572 | 2.348412038 | 15.4 |
| 7.72469E-08 | 0.107084332 | 87 | -1.408349231 | 0.313113278 | 5.424342105 | 2.700323111 | 8.8 |
| 7.74814E-08 | 0.104165302 | 9 | -1.429079279 | 0.318819512 | 5.551155116 | 2.726293753 | 9.1 |
| 7.75143E-08 | 0.19806486 | 69 | -1.045020452 | 0.33264489 | 3.119565217 | 1.290836541 | 44.8 |
| 7.78493E-08 | 0.15580692 | 62 | -1.176273047 | 0.347146339 | 3.754143646 | 1.691340556 | 27.6 |
| 7.81855E-08 | 0.068335055 | 67 | -1.709492144 | 0.271330355 | 9.195519348 | 4.18828767 | 1.8 |
| 7.94475E-08 | 0.141996715 | 54 | -1.232695038 | 0.327331097 | 4.045278137 | 1.959900886 | 22.7 |
| 7.96747E-08 | 0.05179033 | 8 | -1.857153814 | 0.23273021 | 12.50100402 | 5.351841743 | 0.4 |
| 7.98109E-08 | 0.171308257 | 61 | -1.111898946 | 0.351002857 | 3.480406386 | 1.60304284 | 31.1 |
| 7.98632E-08 | 0.015347036 | 60 | -2.086089792 | 0.182644749 | 38.337 | 11.27086791 | 0 |
| 8.01717E-08 | 0.029186794 | 35 | -2.041032616 | 0.183033598 | 21.92592593 | 7.99752708 | 0.1 |
| 8.02683E-08 | 0.077663736 | 31 | -1.637813272 | 0.27513677 | 7.854657114 | 3.883730423 | 2.3 |
| 8.13490E-08 | 0.062851419 | 34 | -1.761976417 | 0.258065989 | 10.15656566 | 4.642174049 | 1 |
| 8.13954E-08 | 0.082687839 | 58 | -1.59664563 | 0.284604228 | 7.356701031 | 3.521800225 | 3 |
| 8.27291E-08 | 0.043498562 | 10 | -1.930263091 | 0.226905057 | 15.218 | 6.20201104 | 0 |
| 8.31551E-08 | 0.065013744 | 32 | -1.733910921 | 0.254109252 | 9.656565657 | 4.405074783 | 1 |
| 8.32055E-08 | 0.198865815 | 7 | -1.057386001 | 0.360799442 | 3.336148649 | 1.514792497 | 40.8 |
| 8.34291E-08 | 0.04059482 | 20 | -1.961650005 | 0.21302687 | 16.272 | 6.307281332 | 0 |
| 8.36063E-08 | 0.105020254 | 73 | -1.464206677 | 0.297449383 | 5.857451404 | 2.875278334 | 7.4 |
| 8.38460E-08 | 0.022189871 | 68 | -2.094246755 | 0.188941813 | 29.35 | 9.70008204 | 0 |
| 8.44043E-08 | 0.173551287 | 94 | -1.120202835 | 0.367050897 | 3.553284672 | 1.739338917 | 31.5 |
| 8.48633E-08 | 0.007450883 | 82 | -2.009825318 | 0.220298767 | 59.352 | 15.85128889 | 0 |
| 8.50673E-08 | 0.038016401 | 81 | -2.002247439 | 0.203001912 | 18.04 | 6.680646004 | 0 |
| 8.52202E-08 | 0.101056905 | 39 | -1.482953891 | 0.304604221 | 6.034151547 | 3.054509618 | 6.3 |
| 8.53627E-08 | 0.068402003 | 95 | -1.743708689 | 0.26522379 | 9.977732794 | 4.706535812 | 1.2 |
| 8.60217E-08 | 0.150985712 | 16 | -1.202945275 | 0.341059198 | 4.100250627 | 2.01000632 | 20.2 |
| 8.70011E-08 | 0.160138506 | 29 | -1.201880229 | 0.326978693 | 3.853887399 | 1.83591088 | 25.4 |
| 8.70292E-08 | 0.060189765 | 80 | -1.806914909 | 0.258185685 | 11.13622603 | 4.816701739 | 0.9 |
| 8.77588E-08 | 0.022669986 | 70 | -2.095047571 | 0.184581303 | 29.638 | 9.70026036 | 0 |
| 8.78073E-08 | 0.048159959 | 26 | -1.908398816 | 0.226948388 | 14.48348348 | 5.882745181 | 0.1 |
| 8.82687E-08 | 0.011916842 | 60 | -2.099889315 | 0.187705653 | 47.196 | 13.15022861 | 0 |
| 8.85433E-08 | 0.003956779 | 87 | -1.906430565 | 0.239726704 | 76.384 | 18.27053246 | 0 |
| 8.87385E-08 | 0.104898223 | 9 | -1.4714337 | 0.306510486 | 6.021164021 | 2.959396254 | 5.5 |
| 8.90021E-08 | 0.055814096 | 5 | -1.855867407 | 0.24928679 | 12.59638554 | 5.346839294 | 0.4 |
| 8.96809E-08 | 0.155377217 | 63 | -1.196451693 | 0.352137991 | 4.075255102 | 1.9556252 | 21.6 |
| 8.99073E-08 | 0.023796868 | 77 | -2.097168783 | 0.180597954 | 29.654 | 9.609090416 | 0 |
| 1.01063E-09 | 0.16210266 | 90 | -1.145825701 | NA | 2 | NA | 99.9 |
| 1.01121E-09 | 0.02783863 | 96 | -1.091958642 | 0.161187493 | 2.046511628 | 0.213082632 | 95.7 |
| 1.04348E-09 | 0.110131915 | 50 | -0.630464825 | NA | 2 | NA | 99.9 |
| 1.04485E-09 | 0.113399701 | 89 | -1.145825701 | NA | 2 | NA | 99.9 |
| 1.07424E-09 | 0.055495772 | 82 | -1.047470297 | 0.16814295 | 2 | 0 | 98.8 |
| 1.16816E-09 | 0.008937843 | 7 | -0.977191086 | 0.301575353 | 2.452755906 | 0.772381643 | 74.6 |
| 1.21365E-09 | 0.125557871 | 61 | -1.071054154 | NA | 2 | NA | 99.9 |
| 1.25418E-09 | 0.09047163 | 49 | -1.02834383 | 0.302137564 | 2.166666667 | 0.389249472 | 98.8 |
| 1.31478E-09 | 0.039668732 | 61 | -1.120522179 | 0.138447973 | 2.125 | 0.336010753 | 96.8 |
| 1.40847E-09 | 0.014239514 | 87 | -1.011873337 | 0.280916267 | 2.387254902 | 0.613505083 | 79.6 |
| 1.51991E-09 | 0.119043647 | 65 | -0.995746611 | 0.07477731 | 2 | 0 | 99.7 |
| 1.52486E-09 | 0.186840658 | 79 | -1.145825701 | 0 | 2 | 0 | 99.8 |
| 1.62228E-09 | 0.013049045 | 11 | -1.064911351 | 0.29785085 | 2.467857143 | 0.7613201 | 72 |
| 1.62990E-09 | 0.090763131 | 87 | -1.145825701 | 0.076379537 | 2 | 0 | 99.7 |
| 1.68352E-09 | 0.127417494 | 44 | -0.84007784 | 0.328940047 | 2 | 0 | 99.3 |
| 1.76480E-09 | 0.043105804 | 9 | -1.11251557 | 0.207585879 | 2.218181818 | 0.497806637 | 94.5 |
| 1.81013E-09 | 0.083598026 | 91 | -1.094679242 | 0.141481647 | 2 | 0 | 98.7 |
| 1.85998E-09 | 0.025107286 | 24 | -1.089896368 | 0.21343108 | 2.291666667 | 0.601571369 | 85.6 |
| 1.88701E-09 | 0.032777442 | 37 | -1.109574203 | 0.195837684 | 2.26744186 | 0.540684068 | 91.4 |
| 1.88956E-09 | 0.022238668 | 56 | -1.088972753 | 0.216306205 | 2.353773585 | 0.625170995 | 78.8 |
| 1.89115E-09 | 0.050529576 | 78 | -1.119976461 | 0.214985229 | 2.125 | 0.40430377 | 96 |
| 1.90273E-09 | 0.106539955 | 82 | -0.920335991 | 0.24306698 | 2 | 0 | 98.7 |
| 1.93148E-09 | 0.021324877 | 89 | -1.092682768 | 0.207997235 | 2.327014218 | 0.570864086 | 78.9 |
| 1.95247E-09 | 0.112849528 | 1 | -1.006942223 | 0.224500291 | 2.090909091 | 0.301511345 | 98.9 |
| 1.96626E-09 | 0.117030422 | 62 | -1.339250771 | 0.123420036 | 3 | 0 | 99.8 |
| 1.96692E-09 | 0.03574858 | 94 | -1.12390884 | 0.170902929 | 2.208791209 | 0.435034699 | 90.9 |
| 2.00049E-09 | 0.024377374 | 57 | -1.087557527 | 0.242330438 | 2.254143646 | 0.539080453 | 81.9 |
| 2.04593E-09 | 0.023275372 | 26 | -1.107615277 | 0.204715163 | 2.292079208 | 0.637810803 | 79.8 |
| 2.11450E-09 | 0.023923197 | 94 | -1.088545706 | 0.227540295 | 2.315508021 | 0.560266002 | 81.3 |
| 2.16334E-09 | 0.006863784 | 73 | -0.991580847 | 0.355518766 | 3.04886562 | 1.331852752 | 42.7 |
| 2.18700E-09 | 0.011080759 | 44 | -1.034890037 | 0.322627879 | 2.699763593 | 0.942505957 | 57.7 |
| 2.20747E-09 | 0.037835737 | 23 | -1.098075992 | 0.232314364 | 2.210526316 | 0.549321751 | 92.4 |
| 2.21115E-09 | 0.013832197 | 29 | -1.085793598 | 0.276607108 | 2.608478803 | 0.817819008 | 59.9 |
| 2.24444E-09 | 0.091031659 | 76 | -1.05031108 | 0.173292801 | 2 | 0 | 98 |
| 2.26867E-09 | 0.036237396 | 77 | -1.125284295 | 0.179306906 | 2.188034188 | 0.434138543 | 88.3 |
| 2.28172E-09 | 0.000889691 | 86 | -0.455335318 | 0.695653761 | 4.59558011 | 2.068511906 | 9.5 |
| 2.28676E-09 | 0.024502224 | 54 | -1.093834287 | 0.191635516 | 2.288770053 | 0.510146601 | 81.3 |
| 2.32057E-09 | 0.042698751 | 70 | -1.107658027 | 0.188258218 | 2.240963855 | 0.508233705 | 91.7 |
| 2.33343E-09 | 0.121074273 | 38 | -1.072394146 | 0.122076717 | 2 | 0 | 99.4 |
| 2.41399E-09 | 0.131678674 | 52 | -1.096961473 | 0.208428635 | 2.222222222 | 0.666666667 | 99.1 |
| 2.47316E-09 | 0.155659682 | 68 | -1.054773253 | 0.162575919 | 2 | 0 | 99.6 |
| 2.50081E-09 | 0.183060031 | 59 | -0.818197687 | 0.054008489 | 2 | 0 | 99.8 |
| 2.50143E-09 | 0.009252218 | 23 | -1.048437569 | 0.332457069 | 2.979202773 | 1.191514 | 42.3 |
| 2.51626E-09 | 0.123918922 | 39 | -1.066741817 | 0.20780973 | 2 | 0 | 98.9 |
| 2.56349E-09 | 0.1252226 | 15 | -0.998694593 | 0.102331873 | 2 | 0 | 99.8 |
| 2.62040E-09 | 0.097424746 | 78 | -1.029598257 | 0.254620082 | 2.071428571 | 0.267261242 | 98.6 |
| 2.64796E-09 | 0.068651967 | 8 | -1.092471575 | 0.217355616 | 2.242424242 | 0.501890366 | 96.7 |
| 2.68883E-09 | 0.112673204 | 79 | -1.053768259 | 0.138673972 | 2 | 0 | 98.4 |
| 2.77947E-09 | 0.185757721 | 46 | -0.929551012 | 0.413875575 | 2 | 0 | 99.8 |
| 2.78982E-09 | 0.04531321 | 50 | -1.145354862 | 0.170725115 | 2.295238095 | 0.553377905 | 89.5 |
| 2.79885E-09 | 0.143077961 | 73 | -1.096033883 | 0.146023087 | 2.125 | 0.353553391 | 99.2 |
| 2.81030E-09 | 0.155736687 | 20 | -0.726140245 | 0.432820529 | 2 | 0 | 99.7 |
| 2.87415E-09 | 0.184450928 | 36 | -1.146227699 | 0.107259298 | 2 | 0 | 99.6 |
| 2.93223E-09 | 0.175651662 | 32 | -0.886805271 | 0.206043994 | 2 | 0 | 99.4 |
| 2.93377E-09 | 0.138910089 | 67 | -1.050273973 | 0.186310394 | 2 | 0 | 98.7 |
| 2.94838E-09 | 0.054018292 | 97 | -1.083275792 | 0.216980851 | 2.129411765 | 0.371234597 | 91.5 |
| 2.95326E-09 | 0.029782522 | 83 | -1.120805092 | 0.199546373 | 2.331730769 | 0.667067187 | 79.2 |
| 3.05530E-09 | 0.063345484 | 66 | -1.10044798 | 0.175957857 | 2.192307692 | 0.486623482 | 94.8 |
| 3.09290E-09 | 0.142877693 | 81 | -0.986419954 | 0.143515861 | 2.25 | 0.46291005 | 99.2 |
| 3.10830E-09 | 0.199706781 | 99 | -0.925263038 | 0.070826145 | 2 | 0 | 99.7 |
| 3.12244E-09 | 0.157430658 | 66 | -0.892969234 | 0.260908084 | 2 | 0 | 99.6 |
| 3.23711E-09 | 0.118960926 | 96 | -1.013065316 | 0.177162792 | 2.125 | 0.341565026 | 98.4 |
| 3.29130E-09 | 0.153624259 | 12 | -0.962514812 | 0.276923124 | 2 | 0 | 99.6 |
| 3.31882E-09 | 0.057128617 | 4 | -1.12450918 | 0.168827861 | 2.2 | 0.496798259 | 90.5 |
| 3.33679E-09 | 0.134054739 | 16 | -0.96682715 | 0.225471047 | 2 | 0 | 98.9 |
| 3.36112E-09 | 0.090815239 | 85 | -1.112066721 | 0.170547596 | 2.205882353 | 0.41042563 | 96.6 |
| 3.40874E-09 | 0.142667866 | 29 | -0.926319127 | 0.238011007 | 2.076923077 | 0.277350098 | 98.7 |
| 3.45303E-09 | 0.116100218 | 51 | -1.044646007 | 0.186339041 | 2 | 0 | 97.4 |
| 3.45651E-09 | 0.07060834 | 37 | -1.071380657 | 0.19324928 | 2.125 | 0.333711906 | 94.4 |
| 3.47480E-09 | 0.036088665 | 87 | -1.151943963 | 0.171793791 | 2.294642857 | 0.578204936 | 77.6 |
| 3.48376E-09 | 0.180788421 | 35 | -1.145825701 | 0.108016977 | 2 | 0 | 99.8 |
| 3.54029E-09 | 0.195300871 | 50 | -0.848921787 | 0.419352563 | 2 | 0 | 99.3 |
| 3.54969E-09 | 0.190918855 | 20 | -1.150676802 | 0.16203972 | 2.222222222 | 0.440958552 | 99.1 |
| 3.56340E-09 | 0.153815397 | 57 | -0.963341718 | 0.266145164 | 2.181818182 | 0.603022689 | 98.9 |
| 3.60466E-09 | 0.010148286 | 50 | -1.111490862 | 0.320856386 | 3.451923077 | 1.463126473 | 27.2 |
| 3.64423E-09 | 0.093226633 | 90 | -1.114331816 | 0.157515526 | 2.060606061 | 0.242305842 | 96.7 |
| 3.68285E-09 | 0.185766621 | 26 | -0.964049503 | 0.220011664 | 2.125 | 0.353553391 | 99.2 |
| 3.71587E-09 | 0.124887435 | 96 | -0.963318807 | 0.391565437 | 2 | 0 | 98.9 |
| 3.74444E-09 | 0.040985564 | 29 | -1.134469376 | 0.17586001 | 2.317204301 | 0.580237352 | 81.4 |
| 3.83984E-09 | 0.156058522 | 28 | -0.941662284 | 0.352897614 | 2.111111111 | 0.333333333 | 99.1 |
| 3.85759E-09 | 0.106453984 | 65 | -1.059422386 | 0.216225713 | 2.041666667 | 0.204124145 | 97.6 |
| 3.86232E-09 | 0.140252604 | 45 | -1.019159908 | 0.226379193 | 2.090909091 | 0.301511345 | 98.9 |
| 3.87286E-09 | 0.145502936 | 1 | -0.999364589 | 0.29886187 | 2 | 0 | 98.2 |
| 3.88632E-09 | 0.032416763 | 76 | -1.150731117 | 0.189422701 | 2.440860215 | 0.741314683 | 72.1 |
| 3.88713E-09 | 0.047017897 | 20 | -1.14801296 | 0.197657482 | 2.372262774 | 0.594178861 | 86.3 |
| 3.90645E-09 | 0.045846424 | 6 | -1.159640492 | 0.173333038 | 2.386503067 | 0.641421583 | 83.7 |
| 3.98044E-09 | 0.117140945 | 68 | -1.019665069 | 0.26025968 | 2.111111111 | 0.323380833 | 98.2 |
| 4.03000E-09 | 0.110095979 | 10 | -1.09189299 | 0.20424468 | 2.161290323 | 0.373878251 | 96.9 |
| 4.04572E-09 | 0.078672153 | 59 | -1.067736191 | 0.187764467 | 2.080645161 | 0.274512211 | 93.8 |
| 4.12910E-09 | 0.054761298 | 85 | -1.129025389 | 0.19543459 | 2.278195489 | 0.56880235 | 86.7 |
| 4.14914E-09 | 0.066314677 | 56 | -1.129088865 | 0.163070482 | 2.23 | 0.489382212 | 90 |
| 4.17794E-09 | 0.0164626 | 75 | -1.164697329 | 0.256826852 | 3.111111111 | 1.251665557 | 37.9 |
| 4.20651E-09 | 0.087631791 | 90 | -1.060111267 | 0.19636677 | 2.140625 | 0.393082547 | 93.6 |
| 4.21081E-09 | 0.064123585 | 21 | -1.091860631 | 0.184766771 | 2.18 | 0.435310176 | 90 |
| 4.21166E-09 | 0.137810585 | 11 | -0.949375807 | 0.27348087 | 2.055555556 | 0.23570226 | 98.2 |
| 4.21670E-09 | 0.081031449 | 55 | -1.103999122 | 0.221934433 | 2.171428571 | 0.416034717 | 93 |
| 4.22175E-09 | 0.000272418 | 2 | -0.690351899 | 0.574334798 | 7.517694641 | 2.974408447 | 1.1 |
| 4.23302E-09 | 0.125799242 | 99 | -1.079104351 | 0.110959711 | 2.19047619 | 0.402373908 | 97.9 |
| 4.24265E-09 | 0.15514505 | 25 | -1.015750775 | 0.249887857 | 2 | 0 | 98.6 |
| 4.24356E-09 | 0.100546756 | 69 | -1.077996209 | 0.173001571 | 2.171428571 | 0.452815654 | 96.5 |
| 4.29833E-09 | 0.024542987 | 87 | -1.183254931 | 0.235510568 | 2.779017857 | 1.01346943 | 55.2 |
| 4.30986E-09 | 0.098298211 | 67 | -1.06413227 | 0.14941474 | 2.054054054 | 0.229243435 | 96.3 |
| 4.34849E-09 | 0.183250244 | 33 | -0.924007656 | 0.360950387 | 2.4 | 0.547722558 | 99.5 |
| 4.39763E-09 | 0.185077495 | 72 | -0.816890897 | 0.325853019 | 2.083333333 | 0.288675135 | 98.8 |
| 4.44240E-09 | 0.08427651 | 70 | -1.10003641 | 0.187711485 | 2.203703704 | 0.450560642 | 94.6 |
| 4.44845E-09 | 0.017813951 | 9 | -1.17632356 | 0.257289987 | 3.014950166 | 1.182840041 | 39.8 |
| 4.45830E-09 | 0.175059041 | 23 | -0.939003085 | 0.225445017 | 2.083333333 | 0.288675135 | 98.8 |
| 4.46223E-09 | 0.01012841 | 35 | -1.174938511 | 0.320436358 | 3.901112485 | 1.695979885 | 19.1 |
| 4.61902E-09 | 0.15531995 | 15 | -0.967296223 | 0.250404309 | 2.066666667 | 0.25819889 | 98.5 |
| 4.62164E-09 | 0.147314766 | 97 | -0.963441291 | 0.298049124 | 2.066666667 | 0.25819889 | 98.5 |
| 4.65305E-09 | 0.025544625 | 33 | -1.158958286 | 0.224263791 | 2.72611465 | 1.001773735 | 52.9 |
| 4.65635E-09 | 0.125795606 | 82 | -1.074312293 | 0.212691774 | 2.129032258 | 0.340777101 | 96.9 |
| 4.71245E-09 | 0.07716111 | 82 | -1.119875884 | 0.19582935 | 2.255813953 | 0.535472227 | 91.4 |
| 4.76454E-09 | 0.18308596 | 26 | -0.912781971 | 0.368044164 | 2 | 0 | 98.6 |
| 4.79342E-09 | 0.189459864 | 1 | -0.910850145 | 0.353987572 | 2.125 | 0.353553391 | 99.2 |
| 4.81383E-09 | 0.045104414 | 9 | -1.156951291 | 0.167607259 | 2.376470588 | 0.669524521 | 74.5 |
| 4.82194E-09 | 0.002499506 | 61 | -0.874957485 | 0.551042377 | 7.516683519 | 3.228945304 | 1.1 |
| 4.91357E-09 | 0.014005405 | 56 | -1.199227029 | 0.3053136 | 3.652116402 | 1.645184268 | 24.4 |
| 4.93431E-09 | 0.005905324 | 98 | -1.17323854 | 0.390895777 | 5.510615711 | 2.454126162 | 5.8 |
| 4.93693E-09 | 0.196083065 | 97 | -0.990675341 | 0.312793344 | 2.125 | 0.353553391 | 99.2 |
| 4.99813E-09 | 0.041186126 | 2 | -1.148684672 | 0.213508447 | 2.413669065 | 0.763395553 | 72.2 |
| 5.07083E-09 | 0.07201194 | 9 | -1.101236057 | 0.252893583 | 2.237623762 | 0.427750274 | 89.9 |
| 5.11498E-09 | 0.125991801 | 68 | -1.09249882 | 0.186480789 | 2.117647059 | 0.409338408 | 96.6 |
| 5.11716E-09 | 0.148427856 | 76 | -0.940659832 | 0.231109065 | 2.076923077 | 0.277350098 | 98.7 |
| 5.17769E-09 | 0.012524331 | 45 | -1.203154545 | 0.30092923 | 3.784747847 | 1.752314048 | 18.7 |
| 5.23359E-09 | 0.092135821 | 18 | -1.072870612 | 0.194604565 | 2.16 | 0.369074811 | 92.5 |
| 5.29038E-09 | 0.102290526 | 95 | -1.040962984 | 0.198739937 | 2.102040816 | 0.367701076 | 95.1 |
| 5.31548E-09 | 0.07914786 | 25 | -1.108323179 | 0.2058043 | 2.232876712 | 0.540589903 | 92.7 |
| 5.33621E-09 | 0.090563766 | 34 | -1.093543323 | 0.18512012 | 2.171052632 | 0.412735844 | 92.4 |
| 5.38098E-09 | 0.157451848 | 38 | -0.98324002 | 0.202145936 | 2 | 0 | 99.1 |
| 5.40690E-09 | 0.1850562 | 14 | -0.800911792 | 0.303986453 | 2 | 0 | 99.1 |
| 5.47559E-09 | 0.011867331 | 90 | -1.251686516 | 0.290839207 | 4.154121864 | 1.934055869 | 16.3 |
| 5.48605E-09 | 0.015551803 | 51 | -1.228041601 | 0.272263023 | 3.557463672 | 1.546779523 | 24.3 |
| 5.50414E-09 | 0.060750666 | 64 | -1.167958256 | 0.172239781 | 2.366459627 | 0.68638037 | 83.9 |
| 5.61863E-09 | 0.035996163 | 63 | -1.183029093 | 0.210397859 | 2.610497238 | 0.899492178 | 63.8 |
| 5.68850E-09 | 0.121377724 | 88 | -1.014565598 | 0.29560012 | 2.261904762 | 0.543678699 | 95.8 |
| 5.70741E-09 | 0.083854371 | 65 | -1.100602574 | 0.195691176 | 2.157894737 | 0.394531488 | 90.5 |
| 5.75030E-09 | 0.01947594 | 38 | -1.240834241 | 0.254003225 | 3.417266187 | 1.466726445 | 30.5 |
| 5.76505E-09 | 0.152226504 | 39 | -0.977476766 | 0.280358507 | 2.047619048 | 0.21821789 | 97.9 |
| 5.89865E-09 | 0.163066009 | 82 | -1.011577341 | 0.266066545 | 2.166666667 | 0.514495755 | 98.2 |
| 5.94085E-09 | 0.130459109 | 32 | -0.999523754 | 0.26285265 | 2.171428571 | 0.452815654 | 96.5 |
| 5.94539E-09 | 0.18194444 | 78 | -0.880514104 | 0.56194322 | 2.066666667 | 0.25819889 | 98.5 |
| 5.99600E-09 | 0.087536784 | 85 | -1.100714669 | 0.196550605 | 2.173469388 | 0.431384972 | 90.2 |
| 6.01210E-09 | 0.021766128 | 64 | -1.224374271 | 0.253383362 | 3.251515152 | 1.332589345 | 34 |
| 6.07143E-09 | 0.154425005 | 98 | -0.976111559 | 0.219431464 | 2.088235294 | 0.287902241 | 96.6 |
| 6.08898E-09 | 0.054544437 | 75 | -1.156228232 | 0.188529885 | 2.422413793 | 0.740315389 | 76.8 |
| 6.10043E-09 | 0.081826466 | 73 | -1.11798194 | 0.235621054 | 2.344537815 | 0.602775709 | 88.1 |
| 6.10749E-09 | 0.116399301 | 41 | -1.003188619 | 0.263561488 | 2.173076923 | 0.382004714 | 94.8 |
| 6.14038E-09 | 0.13420271 | 4 | -0.989565139 | 0.268687092 | 2.088235294 | 0.287902241 | 96.6 |
| 6.17449E-09 | 0.03962968 | 86 | -1.187697909 | 0.226122085 | 2.619302949 | 0.915724372 | 62.7 |
| 6.18262E-09 | 0.161237583 | 10 | -1.012258715 | 0.274510162 | 2.090909091 | 0.294244943 | 97.8 |
| 6.18684E-09 | 0.012842356 | 44 | -1.24529691 | 0.293635728 | 4.223760092 | 1.894623041 | 13.3 |
| 6.19671E-09 | 0.128649514 | 16 | -1.104318441 | 0.162947602 | 2.146341463 | 0.357839043 | 95.9 |
| 6.20313E-09 | 0.072950294 | 19 | -1.109074107 | 0.205247864 | 2.210144928 | 0.504764143 | 86.2 |
| 6.32830E-09 | 0.064723744 | 36 | -1.156063106 | 0.191197798 | 2.411428571 | 0.77451187 | 82.5 |
| 6.34873E-09 | 0.099776515 | 91 | -1.092183265 | 0.180593092 | 2.138297872 | 0.376773622 | 90.6 |
| 6.34945E-09 | 0.165175497 | 93 | -0.854390466 | 0.33572465 | 2.096774194 | 0.300537154 | 96.9 |
| 6.38891E-09 | 0.110258902 | 97 | -1.085744336 | 0.202904705 | 2.15 | 0.40442468 | 94 |
| 6.40015E-09 | 0.050953377 | 14 | -1.172790902 | 0.196916738 | 2.536977492 | 0.79782923 | 68.9 |
| 6.41778E-09 | 0.190737532 | 89 | -0.842880338 | 0.295660345 | 2 | 0 | 99 |
| 6.47170E-09 | 0.060775701 | 82 | -1.163662857 | 0.178143702 | 2.450980392 | 0.757588226 | 79.6 |
| 6.51853E-09 | 0.004617782 | 71 | -1.210709198 | 0.399831 | 7.283076923 | 3.056932534 | 2.5 |
| 6.60017E-09 | 0.025945115 | 24 | -1.23999063 | 0.238003999 | 3.253968254 | 1.357640499 | 37 |
| 6.61228E-09 | 0.155039291 | 80 | -1.077182833 | 0.198121309 | 2.192307692 | 0.491465626 | 97.4 |
| 6.65150E-09 | 0.089377494 | 70 | -1.122540443 | 0.212761949 | 2.292035398 | 0.677197691 | 88.7 |
| 6.74522E-09 | 0.028716439 | 30 | -1.229410947 | 0.225405153 | 3.053872054 | 1.223901989 | 40.6 |
| 6.75761E-09 | 0.181987218 | 95 | -0.90080269 | 0.283784028 | 2.041666667 | 0.204124145 | 97.6 |
| 6.80299E-09 | 0.035593864 | 53 | -1.218523541 | 0.218827111 | 2.816283925 | 1.026666403 | 52.1 |
| 6.80899E-09 | 0.139976064 | 10 | -0.92724971 | 0.31538391 | 2.051282051 | 0.223455865 | 96.1 |
| 6.82377E-09 | 0.026322939 | 33 | -1.227555743 | 0.254452956 | 3.223088924 | 1.422529813 | 35.9 |
| 6.83526E-09 | 0.02185311 | 46 | -1.255990055 | 0.26123279 | 3.510608204 | 1.556613615 | 29.3 |
| 6.88001E-09 | 0.139843747 | 58 | -1.0093959 | 0.250296208 | 2.074074074 | 0.264350529 | 94.6 |
| 6.99125E-09 | 0.112092321 | 9 | -1.05410803 | 0.242560421 | 2.22972973 | 0.454725487 | 92.6 |
| 7.00522E-09 | 0.066422391 | 31 | -1.159830873 | 0.184458943 | 2.422535211 | 0.636810415 | 78.7 |
| 7.06315E-09 | 0.185137758 | 25 | -1.033897579 | 0.244493514 | 2.076923077 | 0.277350098 | 98.7 |
| 7.07613E-09 | 0.041351063 | 42 | -1.205124201 | 0.206045411 | 2.772839506 | 1.072958843 | 59.5 |
| 7.10128E-09 | 0.111405219 | 82 | -1.088357797 | 0.225200353 | 2.222222222 | 0.451046427 | 92.8 |
| 7.11413E-09 | 0.047240915 | 57 | -1.186310723 | 0.200375135 | 2.617283951 | 0.911813098 | 67.6 |
| 7.12264E-09 | 0.095681196 | 32 | -1.087930411 | 0.213895125 | 2.196261682 | 0.422017891 | 89.3 |
| 7.12753E-09 | 0.153702421 | 87 | -0.962651569 | 0.229813026 | 2.136363636 | 0.347141757 | 95.6 |
| 7.18259E-09 | 0.150989175 | 87 | -0.9925314 | 0.162399352 | 2.151515152 | 0.364109541 | 96.7 |
| 7.18950E-09 | 0.186493724 | 90 | -0.768900939 | 0.446322541 | 2.083333333 | 0.282329851 | 97.6 |
| 7.21488E-09 | 0.137496908 | 79 | -1.047597322 | 0.233747406 | 2.156862745 | 0.418212818 | 94.9 |
| 7.27193E-09 | 0.001572966 | 77 | -1.021215784 | 0.525489134 | 10.99098196 | 3.964602509 | 0.2 |
| 7.29832E-09 | 0.111544153 | 12 | -1.04188503 | 0.224923369 | 2.171052632 | 0.379057002 | 92.4 |
| 7.37434E-09 | 0.070972038 | 70 | -1.14641744 | 0.196644062 | 2.435 | 0.684189724 | 80 |
| 7.37955E-09 | 0.179894272 | 10 | -0.921630198 | 0.373420624 | 2.235294118 | 0.562295715 | 98.3 |
| 7.38228E-09 | 0.077346003 | 69 | -1.105137745 | 0.209015308 | 2.307228916 | 0.609676145 | 83.4 |
| 7.38499E-09 | 0.074404424 | 67 | -1.147772967 | 0.214536781 | 2.45508982 | 0.691659318 | 83.3 |
| 7.41888E-09 | 0.184586897 | 15 | -0.988874022 | 0.225523178 | 2.136363636 | 0.351250087 | 97.8 |
| 7.50953E-09 | 0.138095976 | 66 | -1.034205717 | 0.212900905 | 2.127659574 | 0.396562473 | 95.3 |
| 7.51481E-09 | 0.026418274 | 99 | -1.250339013 | 0.251084937 | 3.350591716 | 1.45648474 | 32.4 |
| 7.54150E-09 | 0.077198812 | 13 | -1.157593427 | 0.195277681 | 2.46961326 | 0.792614776 | 81.9 |
| 7.65087E-09 | 0.084947366 | 47 | -1.108905764 | 0.203433845 | 2.298136646 | 0.510449808 | 83.9 |
| 7.65331E-09 | 0.032642781 | 18 | -1.222145963 | 0.254437133 | 2.985790409 | 1.193015202 | 43.7 |
| 7.65991E-09 | 0.025832006 | 8 | -1.265216993 | 0.25145758 | 3.418639053 | 1.568450591 | 32.4 |
| 7.68335E-09 | 0.010311484 | 72 | -1.333162675 | 0.312556897 | 5.847207587 | 2.559182421 | 5.1 |
| 7.68816E-09 | 0.079914311 | 30 | -1.137983494 | 0.206514737 | 2.38547486 | 0.637772786 | 82.1 |
| 7.69386E-09 | 0.188674737 | 90 | -0.989011695 | 0.209983328 | 2 | 0 | 97.7 |
| 7.72114E-09 | 0.162586604 | 75 | -0.943133758 | 0.288029914 | 2.081081081 | 0.276724731 | 96.3 |
| 7.75176E-09 | 0.005815423 | 47 | -1.312571945 | 0.350846263 | 7.816347124 | 3.34008032 | 0.9 |
| 7.80529E-09 | 0.088368467 | 44 | -1.12396216 | 0.194575843 | 2.265151515 | 0.550639441 | 86.8 |
| 7.87197E-09 | 0.10894672 | 7 | -1.063174882 | 0.212754079 | 2.23364486 | 0.506177671 | 89.3 |
| 7.92871E-09 | 0.19843264 | 50 | -0.990612648 | 0.317037141 | 2.166666667 | 0.380693494 | 97.6 |
| 7.99685E-09 | 0.008369269 | 81 | -1.337046159 | 0.338252918 | 6.961934156 | 2.994604067 | 2.8 |
| 8.01327E-09 | 0.193585215 | 25 | -0.900948362 | 0.385229867 | 2.210526316 | 0.418853908 | 98.1 |
| 8.06551E-09 | 0.108890271 | 65 | -1.081055455 | 0.198478403 | 2.165137615 | 0.441247421 | 89.1 |
| 8.14365E-09 | 0.130617238 | 76 | -1.017318034 | 0.252265869 | 2.140350877 | 0.350438322 | 94.3 |
| 8.17114E-09 | 0.075258332 | 23 | -1.138779245 | 0.222082313 | 2.43255814 | 0.732337485 | 78.5 |
| 8.18483E-09 | 0.050371983 | 67 | -1.201021438 | 0.205017868 | 2.640826873 | 0.88901607 | 61.3 |
| 8.24470E-09 | 0.024547216 | 74 | -1.277545786 | 0.253740075 | 3.575635877 | 1.636741586 | 25.3 |
| 8.26690E-09 | 0.092183487 | 54 | -1.143499769 | 0.161789778 | 2.32885906 | 0.608975136 | 85.1 |
| 8.30507E-09 | 0.081704416 | 64 | -1.116203165 | 0.246528133 | 2.424418605 | 0.802107756 | 82.8 |
| 8.31030E-09 | 0.193022601 | 49 | -0.8959442 | 0.297043347 | 2.038461538 | 0.196116135 | 97.4 |
| 8.35460E-09 | 0.188420479 | 37 | -0.843520223 | 0.400404285 | 2.071428571 | 0.262265264 | 97.2 |
| 8.40694E-09 | 0.183921943 | 52 | -0.976813403 | 0.263261996 | 2.071428571 | 0.262265264 | 97.2 |
| 8.42181E-09 | 0.172085151 | 75 | -0.971984654 | 0.284262444 | 2.137931034 | 0.350931203 | 97.1 |
| 8.43248E-09 | 0.17370966 | 50 | -0.973581436 | 0.266502793 | 2.142857143 | 0.355035801 | 96.5 |
| 8.43370E-09 | 0.196662612 | 54 | -0.91649985 | 0.250177244 | 2.096774194 | 0.300537154 | 96.9 |
| 8.44681E-09 | 0.068571065 | 25 | -1.139934404 | 0.197227753 | 2.4921875 | 0.821248153 | 74.4 |
| 8.44749E-09 | 0.189700816 | 93 | -0.998830845 | 0.260769879 | 2.064516129 | 0.249731038 | 96.9 |
| 8.46323E-09 | 0.125887308 | 44 | -1.055804236 | 0.231956951 | 2.160714286 | 0.416774878 | 94.4 |
| 8.47772E-09 | 0.073450666 | 80 | -1.142667633 | 0.210646128 | 2.400921659 | 0.720467363 | 78.3 |
| 8.51053E-09 | 0.050994876 | 17 | -1.191753039 | 0.19497868 | 2.613445378 | 0.828996046 | 64.3 |
| 8.54517E-09 | 0.184751248 | 45 | -0.893953272 | 0.380687338 | 2.130434783 | 0.344350222 | 97.7 |
| 8.60084E-09 | 0.151474097 | 82 | -1.036170237 | 0.231960913 | 2.16 | 0.37032804 | 95 |
| 8.60949E-09 | 0.10508107 | 73 | -1.092185479 | 0.227515237 | 2.290909091 | 0.564149999 | 89 |
| 8.63334E-09 | 0.16381068 | 48 | -0.909971561 | 0.325595286 | 2.022222222 | 0.149071198 | 95.5 |
| 8.64179E-09 | 0.057600032 | 10 | -1.195425726 | 0.209854215 | 2.636094675 | 0.892167775 | 66.2 |
| 8.74248E-09 | 0.00904102 | 46 | -1.355739991 | 0.337972087 | 6.850868233 | 2.959202271 | 2.1 |
| 8.78851E-09 | 0.000620664 | 3 | -0.95636245 | 0.479553828 | 13.70870871 | 4.389849176 | 0.1 |
| 8.88195E-09 | 0.028848465 | 34 | -1.28635734 | 0.239059698 | 3.493562232 | 1.505824903 | 30.1 |
| 8.88317E-09 | 0.037878753 | 24 | -1.256832911 | 0.224262038 | 3.073883162 | 1.254131228 | 41.8 |
| 8.90884E-09 | 0.118572492 | 80 | -1.049948124 | 0.257985408 | 2.279069767 | 0.662458217 | 91.4 |
| 8.92040E-09 | 0.142807314 | 43 | -1.009835488 | 0.27524197 | 2.298245614 | 0.596564937 | 94.3 |
| 8.92560E-09 | 0.109100561 | 5 | -1.0777045 | 0.201814829 | 2.238095238 | 0.480475955 | 87.4 |
| 8.98107E-09 | 0.032088454 | 35 | -1.270939853 | 0.24195228 | 3.34769688 | 1.370739574 | 32.7 |
| 8.98286E-09 | 0.121676149 | 71 | -0.990236976 | 0.278631082 | 2.157894737 | 0.394531488 | 90.5 |
| 8.99266E-09 | 0.064758017 | 15 | -1.167522903 | 0.199654594 | 2.503355705 | 0.783879338 | 70.2 |
| 2.86874E-09 | 0.166334062 | 51 | NA | NA | NA | NA | 100 |
| 4.70737E-09 | 0.108242335 | 51 | NA | NA | NA | NA | 100 |
| 8.12744E-09 | 0.093878481 | 55 | NA | NA | NA | NA | 100 |
| 2.26321E-09 | 0.054840814 | 87 | NA | NA | NA | NA | 100 |
| 2.35190E-09 | 0.164402304 | 46 | NA | NA | NA | NA | 100 |
| 1.27626E-09 | 0.171937101 | 95 | NA | NA | NA | NA | 100 |
| 3.19730E-09 | 0.158838896 | 31 | NA | NA | NA | NA | 100 |
| 3.31992E-09 | 0.072217592 | 43 | NA | NA | NA | NA | 100 |
| 7.12420E-09 | 0.045272388 | 13 | NA | NA | NA | NA | 100 |
| 8.33616E-09 | 0.109313414 | 88 | NA | NA | NA | NA | 100 |
| 2.45418E-09 | 0.18871725 | 51 | NA | NA | NA | NA | 100 |
|  |  |  | *NA = Could not be estimated, all replicates were invariable |  |  |  |  |

